# Synteny – a high throughput web tool to streamline causal gene prioritisation and provide insight into protein function

**DOI:** 10.1101/2025.03.16.643559

**Authors:** Harry B. Cutler, Jacqueline Stöckli, Søren Madsen, Stewart W.C. Masson, Oliver K. Fuller, Tom Buckingham Shum, David E. James

## Abstract

Accounting for human genetic evidence can improve the outcomes and impact of basic research studies. However, current approaches are incompatible with the high volume of disease-associated genes that require mechanistic interrogation. *Synteny* responds to an urgent need in systems biology, scaling human genetic analysis to match the throughput of modern omics technologies. This approach prioritises candidates with the strongest human disease relevance and unearths functionally important proteins (https://bigproteomics.shinyapps.io/Synteny/).

## Introduction

Advances in omics-technologies have revolutionised the study and understanding of human disease. However, the complexity of the data generated by these techniques can obscure key biological insights.

A major challenge is that omics studies frequently identify hundreds of disease-associated genes, many of which remain poorly characterised. Even a comprehensive review of the literature often provides insight into just a small subset of these genes^1–3^. Combined with the fact that data analysis approaches like differential expression offer little insight into whether changes are causes or consequences of disease processes, researchers are often drawn to genes with known functions^3,4^. This ‘streetlight’ bias – searching for answers where the light is brightest rather than venturing into the relative unknown – impedes new breakthroughs and signals an unmet need for robust and unbiased tools to prioritise genes with the strongest causal links to human disease.

One approach that is becoming popular is to integrate omics data with findings from human genetics, since genetic variants are causal by definition. Genetic and phenotypic data are now available across large human cohorts and several studies using model systems have demonstrated success in applying such an integrated approach^5–7^. For example, to pinpoint causal genes contained within human GWAS loci, Li et al^6^ integrated genetic associations from a GWAS of human lipid metabolism with gene co-expression networks from mouse liver samples. This identified 54 genes with conserved function in mice and humans, of which 25 had not previously been linked to lipid metabolism. It also motivated experiments to validate the role of Sestrin 1 in regulating cholesterol biosynthesis, identifying a previously unknown therapeutic target. Examples like this emphasise the importance of bridging the gap between studies in humans and model systems. However, the complexity of current approaches provides a considerable challenge to those without specialist bioinformatic training. For example, Keller et al^5^ devised an approach to determine whether loci identified in their mouse GWAS of pancreatic islet function were conserved in humans by mapping syntenic regions between mouse and human GWAS and then testing for overlap with human GWAS loci. Although very rigorous, this approach is technically challenging and only accessible to those with access to genetically diverse mouse populations.

To address these challenges, we developed *Synteny*, a user-friendly web tool that enables anybody with proteomic or transcriptomic data to leverage large-scale human genetic datasets to rank genes by human disease relevance. In contrast to current gene-prioritisation platforms that are limited to searching genes one-at-a-time, such as OpenTargets Genetics^8^ and the Type 2 Diabetes Knowledge Portal^9^, *Synteny* was built to perform analyses at scale – data for several hundred genes can be acquired in minutes. *Synteny* automatically maps human orthologs from 8 common model organisms using the Alliance for Genome Resources database^10^, before acquiring humans phenome-wide association statistics from the Association to Function (A2F) database (https://a2f.hugeamp.org/). Cross-species orthologs are typically highly conserved in terms of structure and function^11^ and so this approach enables findings in model organisms to be leveraged to infer potential human disease mechanisms and gene function, and vice-versa. As of June 2025, A2F reports summary statistics for 898 genetic studies and 1175 phenotypes, enabling unbiased assessment of gene function.

### Synteny powers improved access to human genetic evidence

While technical barriers once limited the ability to integrate human genetic data into broader biological analyses, recent approaches such as the Human Genetic Evidence (HuGE) framework by Dornbos et al ^12^ have greatly expanded access to these insights. The main benefit of HuGE scores is that they provide a single interpretable measure of the strength of association between every gene and every phenotype. They are calculated using a Bayesian model to account for the effects of rare and common genetic variation, the proximity of a gene to the peak genetic association in genome wide association studies and the impact of coding mutations detected in whole exome sequencing^12^. By aggregating genetic data across many independent studies, HuGE scores provide an unbiased and robust assessment of how strongly a gene of interest might relate to human physiology. For example, HuGE scores for peroxisome proliferator activated receptor gamma (*PPARG*), a master-regulator of adipogenesis, show strong associations (HuGE ≥ 30) to body fat percentage and muscle fat infiltration (Fig 1A), consistent with its established role in adipogenesis. While databases like A2F have made it straightforward to access HuGE scores for individual genes, the restriction on searching genes one-at-a-time is incompatible with high-throughput, hypothesis-generating studies where hundreds of disease-associated genes can be identified in a single experiment (Fig 3B).

**Figure 1.**
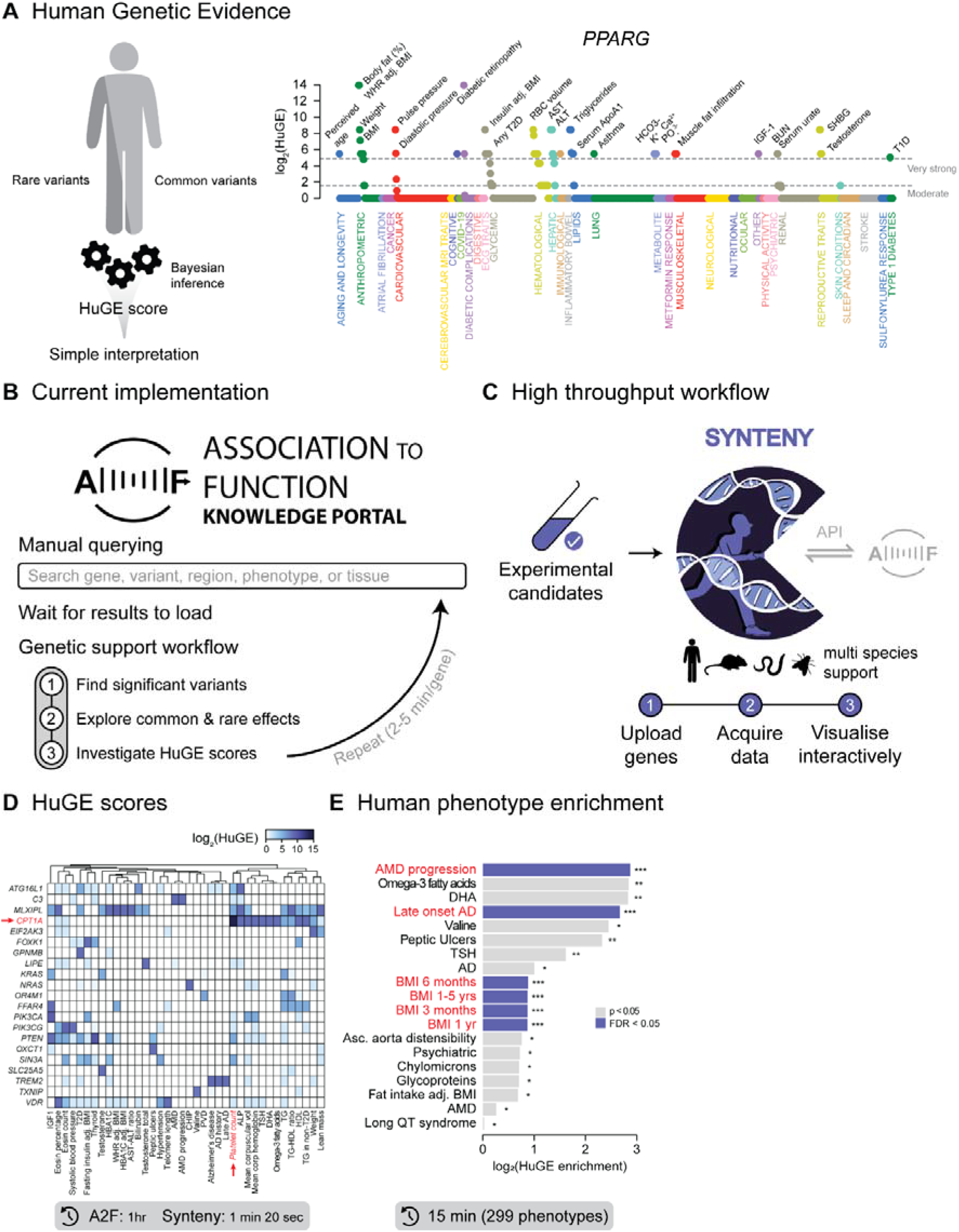
Synteny powers improved access to human genetic evidence. **(A)** Human genetic evidence (HuGE) scores aggregate genetic data from many studies to summarise the strength of association between genes and phenotypes^12^, enabling gene function to be rapidly assessed. Example of HuGE score plot linking *PPARG* to 1175 phenotypes. **(B)** Schematic of the current implementation of the HuGE score framework via the Association to Function web portal. **(C)** Schematic illustrating the high throughput implementation in *Synteny*. API = application programming interface. Examples of data visualisation and analysis in *Synteny* for 52 genes hypothesised to be linked to glucose metabolism^12^ **(D & E). (D)** Heatmap showing associations between genes and human phenotypes, filtered for genes with HuGE scores ≥ 30. **(E)** Results of HPE. Asterisks represent unadjusted p values.

*Synteny* overcomes this limitation by enabling bulk querying of the A2F database (Fig 1C), streamlining the process of integrating human genetic evidence with data from model systems. Its operation is simple, requiring only uploading a list of genes. All results can be exported for analysis in external software and data visualisation plots can be downloaded as high-resolution vector PDFs.

To demonstrate the efficiency of this approach, we used *Synteny* to analyse a list of 52 genes that Dornbos et al^12^ previously used to assess the use of human genetic evidence by metabolism researchers working with model systems. Dornbos et al^12^ selected these genes from studies published in high profile journals between January 2017 and October 2020 that contained any of “diabetes,” “glucose,” or “insulin” in their abstracts. Examining the HuGE scores for these genes, Dornbos et al^12^ identified a striking lack of genetic evidence to support therapeutic targeting of these genes, highlighting an essential need for increased use of human genetic data. However, without efficient tools to query human genetic databases, it remains challenging to ensure that research is targeted towards genes with the strongest links to human disease.

While manually querying each gene using the A2F database required more than an hour, using *Synteny* this process was completed in just 80 seconds (Fig 1D). This boost in efficiency makes this approach much more accessible than existing alternatives, enhancing the potential for increased translatability.

Investigation of HuGE scores across genes frequently reveals well-studied associations, however, the true value of this analysis lies in highlighting relationships that would otherwise have gone unnoticed. For example, while the link between phosphatase and tensin homolog (*PTEN*) and systemic glucose homeostasis is well-established^13,14^, we were surprised by the association between carnitine palmitoyltransferase I (*CPT1A*) and platelet number. CPT1A plays a central role in transporting fatty acids into the mitochondria for oxidation^15,16^ and, consistent with human genetic evidence, recent studies have revealed that the contribution of fatty acids to ATP production in platelets is likely greater than currently appreciated^17,18^. These insights suggest new avenues for exploring platelet pathology and showcase the power of unbiased analyses.

The Human Phenotype Enrichment (HPE) tool in *Synteny* enables users to quantify the extent to which gene sets of interest are supported by human genetic evidence. Similar to existing over-representation analyses for molecular functions, HPE works by comparing HuGE scores for a user-supplied gene set against those obtained by randomly sampling the genome (or a user-supplied background gene set). Consistent with the findings of Dornbos et al^12^, that their list of 52 genes was overall not supported by human genetic evidence, HPE using *Synteny* failed to detect significant enrichment for glycaemic traits that would be expected if these genes were supported by human genetic evidence (Fig 1E). Instead, the 52 genes were enriched for early life Body Mass Index (BMI), Alzheimer’s disease (AD) and age-related macular degeneration (AMD). This outcome supports the conclusions of Dornbos et al^12^, emphasising the importance of interrogating results in the context of human genetics to maximise the extent to which advances in basic research can be translated into human therapeutic breakthroughs. Many existing approaches using human genetic data are designed to test single phenotypes and so here we report both unadjusted and FDR-corrected p values to provide compatibility with these approaches.

To evaluate the sensitivity and robustness of HPE, we performed a series of benchmarking experiments (Fig 2). To determine the degree of enrichment required to detect statistical significance, power analyses were performed using gene sets of varying size between 10-300 genes, obtained by weighted sampling of the HuGE score distribution (Fig 2A: i). Using obesity as an example trait, 90% power was obtained at 4-fold enrichment with 100 genes (Fig 2A: ii). To assess sensitivity more broadly, we extended this analysis to 15 randomly sampled human traits from 16 trait categories (240 phenotypes in total). Except for haematological traits that required a median 40-fold enrichment to reach 80% power, the other 225 traits were broadly consistent, requiring a median enrichment of 1.66-fold (Fig 2B). These results suggest that HPE is highly sensitive for phenome-wide enrichment analysis.

**Figure 2.**
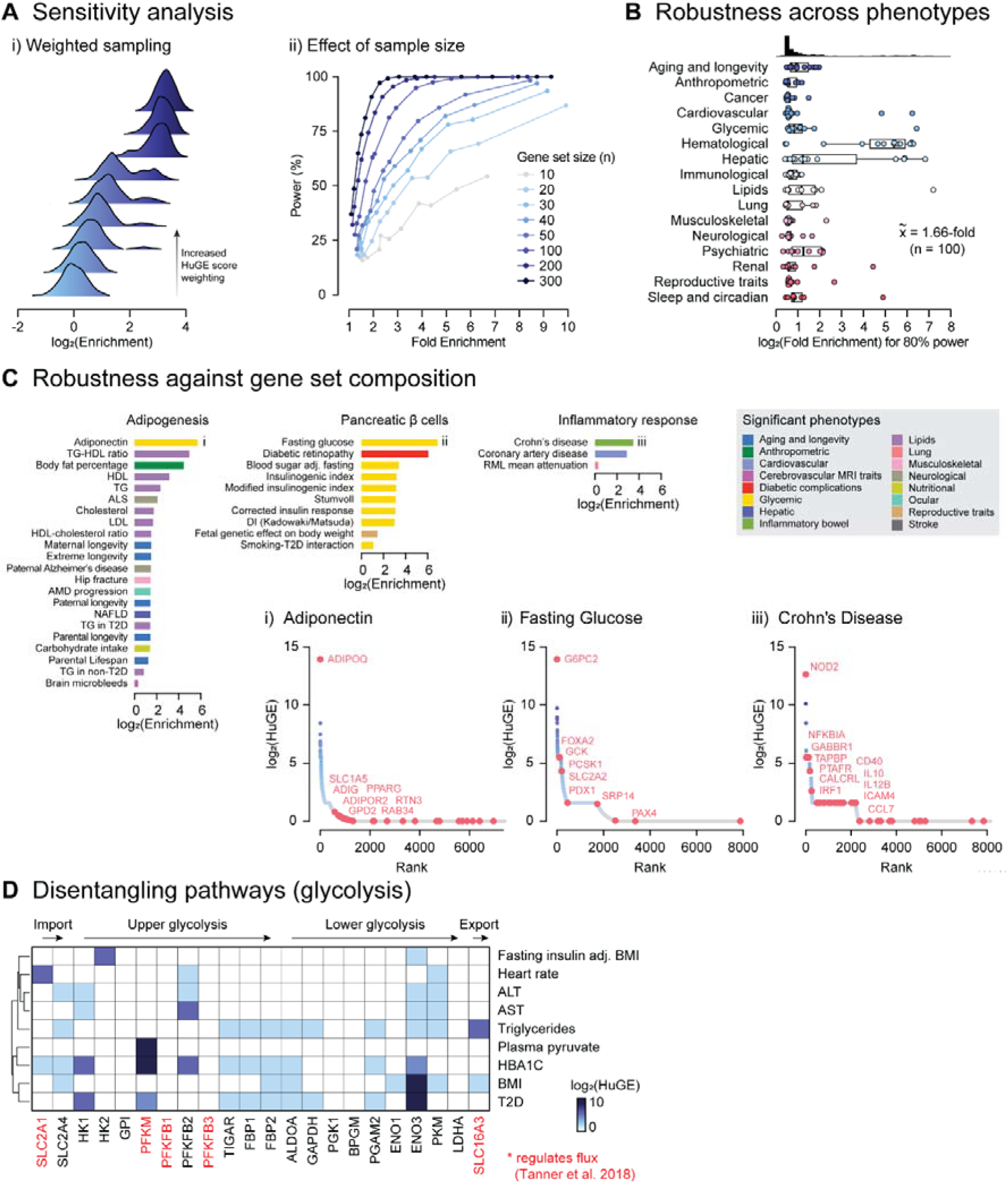
Benchmarking for human phenotype enrichment. **(A)** Example of sensitivity analysis performed using Obesity as a model trait. Random weighted sampling (i) was used to generate enriched gene sets with different numbers of genes that were then subjected to HPE (ii). **(B)** Power simulations for 15 randomly sampled phenotypes from 16 trait categories, with gene sets of size n = 100. Each dot is the average of 1000 permutations per phenotype, showing the degree of enrichment required to reach 80% power. Median enrichment for 80% power (x = 1.66-fold. **(C)** HPE using gene sets from HALLMARK pathways. Genes contributing to the top enrichments for ‘Adipogenesis’, ‘Pancreatic β cells’ and ‘Inflammatory response’ are shown in i-iii, respectively. **(D)** Heatmap presenting HuGE scores for glycolytic enzymes expressed in skeletal muscle for selected phenotypes. Enzymes that regulate total glycolytic flux^20^ are highlighted in red.

To assess the robustness of HPE against varying gene set compositions, we used HPE to analyse several systematically curated HALLMARK pathways (Fig 2C). This identified enrichments that are consistent with known phenotypic associations. For example, the ‘Adipogenesis’ pathway was enriched for adiponectin (an adipokine), body fat percentage and plasma triglyceride (TG) levels; the ‘Pancreatic β cells’ pathway was enriched for fasting glucose and traits related to insulin secretion, a key function of β cells; and the ‘Inflammatory response’ pathway was enriched for Crohn’s disease and coronary artery disease, both well-studied inflammatory conditions. These results suggest that HPE can infer true biological meaning from user-supplied gene sets.

HPE aggregates enrichment signals over entire gene sets, and it is therefore important to determine whether findings are driven by a small number of genes or are distributed across entire the gene set. *Synteny* facilitates this by automatically visualising the contributions of user-supplied genes to an enriched phenotype. For example, while the enrichment for adiponectin in the ‘Adipogenesis’ gene set seems to be primarily driven by the inclusion of the adiponectin (ADIPOQ) gene, the Crohn’s disease enrichment for the ‘Inflammatory response’ pathway is more evenly distributed.

Enabling users to determine which genes contribute most strongly to human traits also addresses a core challenge in omics analysis. Genes belonging to the same pathway are often co-regulated^19^, and this can obscure which genes causally influence the phenotype. For example, although omics analysis might identify several glycolytic enzymes as hits, it may be unclear which gene/s should be pursued experimentally. This is important since, while there are 12 steps in glycolysis – from glucose import to lactate export – only 4 of these appear to regulate total flux through the pathway^20^. Selecting genes that do not regulate flux might result in a false negative result. *Synteny* addresses this challenge by providing access to the effects of human genetic variation. Because the effects of genetic variation are independent of pathway membership, this enables users to disentangle the contributions of individual genes to human disease. Applied to the analysis of glycolytic genes expressed in skeletal muscle, *Synteny* identified very strong genetic evidence (HuGE ≥ 30) for several enzymes that Tanner et al^20^ previously showed to regulate total glycolytic flux (Fig 2D). Thus, *Synteny* provides a powerful tool to assist users in identifying high-priority genes for mechanistic follow-up.

### Synteny leverages human genetic data for discovery

Less than half of all human genes have been studied mechanistically^4,21,22^. To evaluate whether *Synteny* could be used to infer protein function, we applied it to the solute carrier (SLC) superfamily, containing over 400 genes. We chose this family since, despite its members playing crucial roles in many aspects of biology, the substrates and biological functions of many of these proteins remain unknown^23^.

Consistent with their roles as transporters, HPE analysis identified enrichment of the SLC family in regulating circulating levels of a variety of metabolites including creatinine, urate and sodium (Fig 3A). Several human diseases related to dysregulated ion transport, including chronic kidney disease and epilepsy with sclerosis were also enriched.

**Figure 3.**
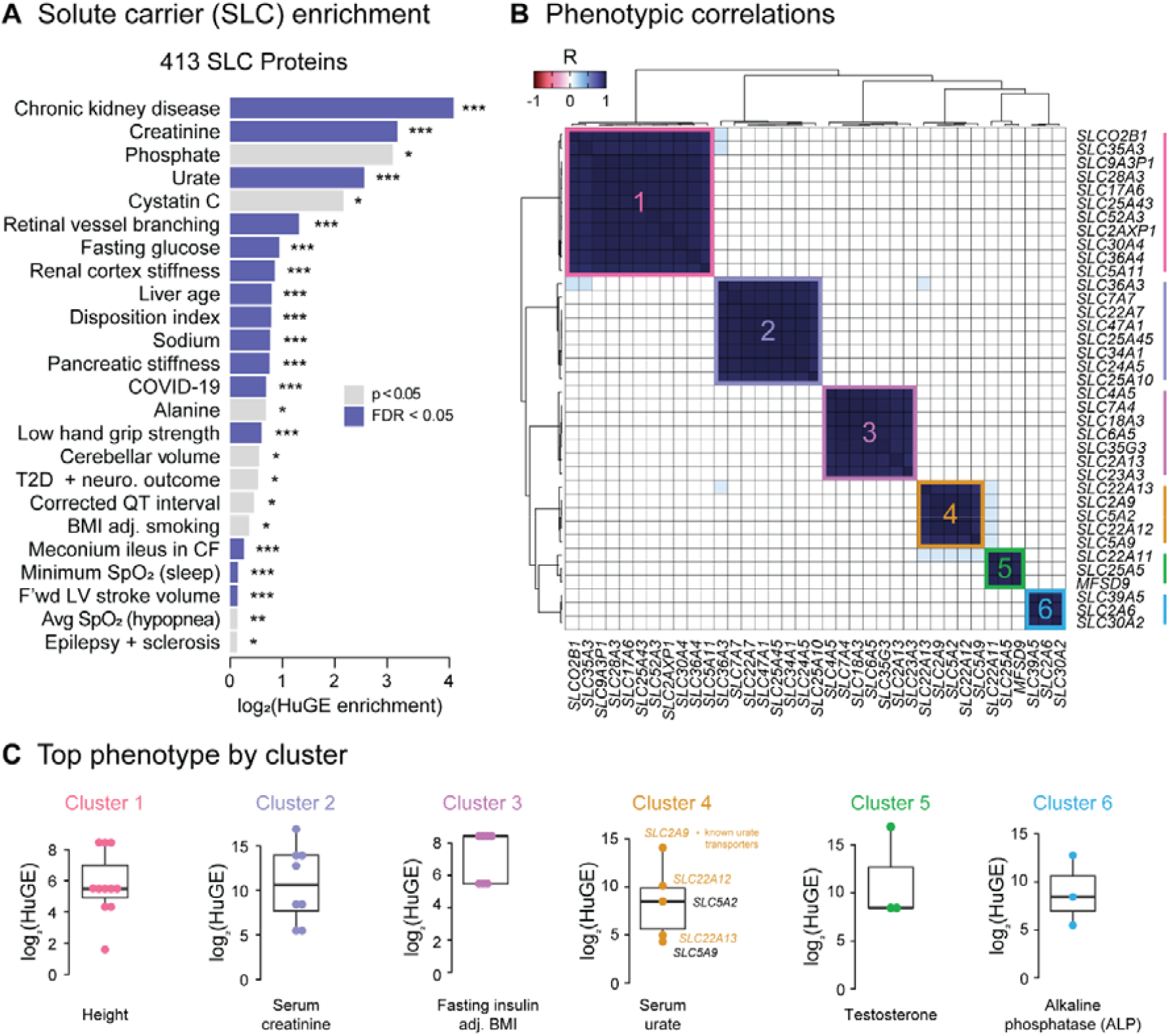
Synteny leverages human genetic data for discovery. **(A)** Results of HPE for 413 members of the solute carrier (SLC) superfamily, only possible using Synteny. Asterisks represent unadjusted p values. **(B)** Correlation heatmap of HuGE scores for SLC genes with more than one R > 0.95 p < 0.01. Clusters contain SLC genes with similar phenotypic associations. **(C)** Top scoring phenotype for each SLC cluster.

To identify putative functions for orphan SLC members – SLCs without known substrates – we identified SLCs with similar associations to human traits by correlating HuGE scores across genes. Such guilt-by-association strategies are implemented in several established tools including WGCNA^24^, GeneBridge^22^ and QENIE^25^, all of which assign putative functions to genes by means of correlations or co-expression networks. This identified 6 clusters containing more than 2 SLC proteins (Fig 3B). Each SLC cluster was enriched for at least one human trait (Fig 3C). Strikingly, cluster 4 was enriched for genes associating with serum urate levels and 3 of the 5 genes in this cluster (*SLC2A9, SLC22A12, SLC22A13*) have previously been shown to actively transport urate^26–28^. This raised the intriguing possibility that the remaining two 2 SLC members in this cluster (*SLC5A2* and *SLC5A9*) may also function as urate transporters.

*SLC5A2* encodes the SGLT2 protein, a sodium-dependent glucose transporter highly expressed in the kidney and a well-established target for Type 2 Diabetes^29,30^. Both genetic and pharmacological evidence support a potential role of *SLC5A2* in urate transport, although this has not been directly tested yet. Specifically, a coding variant in *SLC5A2* has been linked to uricosuria^31^, a condition with high urate levels in the urine, and pharmacological inhibition of SGLT2 decreases the re-absorption of urate from the urine^32^. While it has been suggested that this effect is mediated indirectly through the effects of elevated urinary glucose on urate transport by GLUT9 (*SLC2A9*)^31^ and URAT1 (*SLC22A12*) ^31,33^, studies have shown that GLUT9 is dispensable for the hyperuricosuria^32^ and the impact of glucose on URAT1 remains unclear. This raises the possibility that SGLT2 may directly mediate urate transport, a hypothesis worthy of future investigation.

*SLC5A9* encodes the SGLT4 protein, another sodium-dependent sugar transporter expressed in the small intestine^34,35^, although little else is known about its function. Combined with the fact that approximately 30% of uric acid is excreted into the small intestine, the known overlap between urate and sugar transporters^26^ supports the possibility that *SLC5A9* could also transport urate. Enhancing intestinal urate disposal has been proposed as a therapeutic target for chronic kidney disease^36^ and so it will be important for future studies to clarify whether this might be possible via targeting *SLC5A9*.

These examples highlight how human genetic data can be leveraged to gain insights into protein function and emphasise the utility of *Synteny* in informing targeted validation experiments

## Discussion

Combined with existing bioinformatic tools for identifying genes of interest, *Synteny* provides a systematic and scalable pipeline to maximise the biological insight that can be drawn from omics studies (https://bigproteomics.shinyapps.io/Synteny/).

*Synteny* is not intended to replace existing knowledge portals. These databases contain vast amounts of information and facilitate targeted inspection of individual genes. Rather, *Synteny* complements these resources by helping researchers prioritise where to look, highlighting which genes or pathways are most strongly supported by human genetic evidence. To streamline downstream analysis, *Synteny* automatically generates hyperlinks to the A2F database for genes and phenotypes of interest. This functionality enables users to transition seamlessly from broad scale discovery to targeted experimental follow-up.

Several new analyses are possible using *Synteny*. By searching for genes with shared disease associations, it is possible to assign putative functions to proteins. In an analysis of the SLC family, this approach led us to propose SGLT2 and SGLT4 as putative urate transporters. Combined with the observation that both genetic and pharmacological manipulation of SGLT2 influences plasma urate levels^31,32^, the fact that these genes clustered together with three known urate transporters provides considerable support for the use of human genetic evidence to group genes by their physiological functions.

Beyond functional annotation, *Synteny’s* phenotype enrichment capabilities offer clinical applications. For rare diseases, HPE might reveal related phenotypes that could serve to improve diagnosis or treatment, highlighting the importance of comprehensive phenotype analysis.

As with all tools, however, *Synteny* is not without limitation. Firstly, analysis of genes from diverse tissues could limit mechanistic insight in cases where gene function is not well characterised. To mitigate this, we suggest users stratify genes by tissue expression prior to HPE. This issue does not apply to many common situations where genes sets are obtained by analysing omics data in a specific tissue. Secondly, *Synteny* does not currently incorporate information from Mendelian randomisation or protein quantitative trait loci (pQTL) studies linking proteins to disease. These datasets offer an additional form of evidence to aid candidate prioritisation. We encourage users to make use of data from proteogenomics consortia such as SCALLOP^37^ and the UK BioBank^38^ to perform these analyses once they have identified high-priority candidates using *Synteny*. Finally, *Synteny* does not currently support custom trait-gene mapping using local GWAS data.

*Synteny* addresses a fundamental challenge in modern biology by bridging the gap between studies in model systems and human genetic studies. While the benefits to those working in model systems were discussed above, this approach also offers considerable benefits to those working with human genetic data. Human genome-wide association studies (GWAS) have identified thousands of disease-associated loci. However, these loci often contain many genes and so pinpointing the causal gene/s within these loci remains a persistent challenge. By enabling users to cross-reference GWAS candidates with functional evidence from model systems, *Synteny* helps to prioritise which genes within complex loci are most likely to be causal and merit experimental validation.

## Resource availability

*Synteny* is freely available at the following address: https://bigproteomics.shinyapps.io/Synteny/. Code is open-source and available in the supplementary materials. This can be run locally using RStudio.

## Acknowledgements

This work was supported by an Australian Research Council Laureate Fellowship (to DEJ) and an Australian Government Research Training Program Scholarship (to HBC). The content is solely the responsibility of the authors and does not necessarily represent the official views of the ARC.

We would like to thank several colleagues for their constructive feedback: Marcus Seldin for suggestions relating to enabling broader use of the platform by incorporating the Alliance for Genome Resources database; Alice Williamson for suggestions relating to heatmap visualisation settings and improving the documentation for HuGE correlation analysis; Rob Williams for suggestions relating to the inclusion of metadata in exported results; and Mark Keller for suggestions relating to using *Synteny* with large (> 500) numbers of genes. These conversations greatly improved the user experience of *Synteny*. We welcome future suggestions from the community.

## Author contributions

Conceptualisation, HBC, JS, SM, SWCM, OKF, DEJ; Methodology, HBC; Formal Analysis, HBC; Data curation, HBC; Writing – Original Draft, HBC; Writing – Review & Editing, HBC, JS, SM, SWCM, OKF, DEJ; Visualisation, HBC, TBS; Supervision, DEJ; Funding Acquisition, DEJ.

## Supplemental information

Supplementary file 1: zip file containing source code required to run *Synteny* locally.

## Methods

### Statistical analyses

All analysis and data visualisation were conducted using the R programming environment^39^. Unless otherwise stated, correlations were assessed by Pearson correlation analysis. Significance is represented, with a p value < 0.05 by *, < 0.01 by **, < 0.001 by *** and < 0.0001 by ****.

### Human phenotype enrichment

Human genetic evidence (HuGE) scores were downloaded for each gene. To assess enrichment for a particular phenotype, the same number of genes were randomly selected from the Association to Function database (for naïve enrichment) or randomly from a set of background genes (for background-corrected enrichment). This was repeated for 10,000 permutations to enable the average HuGE score in the user-defined gene set to be compared to the sampled distribution/s. An enrichment score was calculated as the mean fold change of the average scores, and significance was determined by calculating the proportion of sampled scores exceeding that of the user-defined set. Multiple corrections were performed using a false-discovery rate (FDR) correction to account for the number of phenotype tests.

### Benchmarking

#### Sensitivity analysis

Gene sets with progressively greater enrichment were generated by weighted sampling. We performed permutation testing to determine the power of HPE to detect enrichment for gene sets with different levels of enrichment (n = 1000). Power is reported as the proportion of gene sets where enrichment was detected with p < 0.05. Phenotypes in Figure 2B were obtained by randomly sampling 15 phenotypes per trait category.

#### HALLMARK analysis

Human HALLMARK gene sets were downloaded from https://www.gsea-msigdb.org/gsea/msigdb/human/genesets.jsp?collection=H and analysed using HPE. Significantly enriched phenotypes are reported.

### Synteny

#### Web server implementation

*Synteny* is implemented as a Shiny application using the R programming language and is hosted on a shinyapps.io server.

#### Privacy and security

Gene lists are only uploaded if the user accesses *Synteny* via the web server. No user data are retained following session termination. Secure HTTPS connections are used to transfer data to and from the server. The country in which each user is located is logged using a google analytics cookie. This data is used solely to monitor global adoption of the tool and ensure server requirements are sufficient to meet demand. No other data is recorded.

#### Local installation

The code required to run *Synteny* locally is available in the supplementary materials. After installing the required R packages, *Synteny* will still require an internet connection to connect to the Association to Function application programming interface. No data will be uploaded to the *Synteny* shinyapps.io server if running locally.

#### Packages and databases

*Synteny* maps gene names from any organism to human orthologs using the Alliance for Genome Resources database^10^. Human genetic evidence (HuGE) scores are retrieved via an application programming interface connected to the Association to Function database using the jsonlite package^40^. Static heatmaps are created using the ComplexHeatmap package^41^ and interactive plots are created using a combination of ggplot2^42^ and plotly^43^.

#### Exporting results

Results and figures can be explored and visualised within *Synteny* itself or exported for analysis in the users’ software of choice. Data is exportable as excel sheets (.xlsx) and figures are exportable as full resolution vector PDFs.

## References

1. Müller, J.B., Geyer, P.E., Colaço, A.R., Treit, P. V., Strauss, M.T., Oroshi, M., Doll, S., Virreira Winter, S., Bader, J.M., Köhler, N., et al. (2020). The proteome landscape of the kingdoms of life. Nature 2020 582:7813 582, 592–596. 10.1038/s41586-020-2402-x.

2. Hutchison, C.A., Chuang, R.Y., Noskov, V.N., Assad-Garcia, N., Deerinck, T.J., Ellisman, M.H., Gill, J., Kannan, K., Karas, B.J., Ma, L., et al. (2016). Design and synthesis of a minimal bacterial genome. Science (1979) 351. 10.1126/SCIENCE.AAD6253/SUPPL_FILE/AAD6253-HUTCHISONSM.PDF.

3. Kustatscher, G., Collins, T., Gingras, A.C., Guo, T., Hermjakob, H., Ideker, T., Lilley, K.S., Lundberg, E., Marcotte, E.M., Ralser, M., et al. (2022). Understudied proteins: opportunities and challenges for functional proteomics. Nat Methods 19, 774–779. 10.1038/S41592-022-01454-X.

4. Edwards, A.M., Isserlin, R., Bader, G.D., Frye, S. V., Willson, T.M., and Yu, F.H. (2011). Too many roads not taken. Nature 2011 470:7333 470, 163–165. 10.1038/470163a.

5. Keller, M.P., Rabaglia, M.E., Schueler, K.L., Stapleton, D.S., Gatti, D.M., Vincent, M., Mitok, K.A., Wang, Z., Ishimura, T., Simonett, S.P., et al. (2019). Gene loci associated with insulin secretion in islets from nondiabetic mice. J Clin Invest 129, 4419–4432. 10.1172/JCI129143.

6. Li, Z., Votava, J.A., Zajac, G.J.M., Lamming, D.W., Liang, C.-, Yen, E., and Parks Correspondence, B.W. (2020). Integrating Mouse and Human Genetic Data to Move beyond GWAS and Identify Causal Genes in Cholesterol Metabolism. Cell Metab 31, 741-754.e5. 10.1016/j.cmet.2020.02.015.

7. Parker, B.L., Calkin, A.C., Seldin, M.M., Keating, M.F., Tarling, E.J., Yang, P., Moody, S.C., Liu, Y., Zerenturk, E.J., Needham, E.J., et al. (2019). An integrative systems genetic analysis of mammalian lipid metabolism. Nature 567, 187. 10.1038/S41586-019-0984-Y.

8. Mountjoy, E., Schmidt, E.M., Carmona, M., Schwartzentruber, J., Peat, G., Miranda, A., Fumis, L., Hayhurst, J., Buniello, A., Karim, M.A., et al. (2021). An open approach to systematically prioritize causal variants and genes at all published human GWAS trait-associated loci. Nat Genet 53, 1527–1533. 10.1038/S41588-02100945-5.

9. Costanzo, M.C., von Grotthuss, M., Massung, J., Jang, D., Caulkins, L., Koesterer, R., Gilbert, C., Welch, R.P., Kudtarkar, P., Hoang, Q., et al. (2023). The Type 2 Diabetes Knowledge Portal: An open access genetic resource dedicated to type 2 diabetes and related traits. Cell Metab 35, 695-710.e6. 10.1016/J.CMET.2023.03.001.

10. Bult, C.J., and Sternberg, P.W. (2023). The alliance of genome resources: transforming comparative genomics. Mammalian Genome 34, 531. 10.1007/S00335-023-10015-2.

11. Pers, T.H., Karjalainen, J.M., Chan, Y., Westra, H.J., Wood, A.R., Yang, J., Lui, J.C., Vedantam, S., Gustafsson, S., Esko, T., et al. (2015). Biological interpretation of genome-wide association studies using predicted gene functions. Nature Communications 2015 6:1 6, 1–9. 10.1038/ncomms6890.

12. Dornbos, P., Singh, P., Jang, D.K., Mahajan, A., Biddinger, S.B., Rotter, J.I., McCarthy, M.I., and Flannick, J. (2022). Evaluating human genetic support for hypothesized metabolic disease genes. Cell Metab 34, 661–666. 10.1016/j.cmet.2022.03.011.

13. Pal, A., Barber, T.M., Van de Bunt, M., Rudge, S.A., Zhang, Q., Lachlan, K.L., Cooper, N.S., Linden, H., Levy, J.C., Wakelam, M.J.O., et al. (2012). PTEN Mutations as a Cause of Constitutive Insulin Sensitivity and Obesity. New England Journal of Medicine 367, 1002–1011. 10.1056/NEJMOA1113966/SUPPL_FILE/NEJMOA1113966_DISCLOSURES.PDF.

14. Wijesekara, N., Konrad, D., Eweida, M., Jefferies, C., Liadis, N., Giacca, A., Crackower, M., Suzuki, A., Mak, T.W., Kahn, C.R., et al. (2005). Muscle-specific Pten deletion protects against insulin resistance and diabetes. Mol Cell Biol 25, 1135–1145. 10.1128/MCB.25.3.1135-1145.2005.

15. McGarry, J.D., Leatherman, G.F., and Foster, D.W. (1978). Carnitine palmitoyltransferase I. The site of inhibition of hepatic fatty acid oxidation by malonyl-CoA. Journal of Biological Chemistry 253, 4128–4136. 10.1016/S00219258(17)34693-8.

16. Schlaepfer, I.R., and Joshi, M. (2020). CPT1A-mediated Fat Oxidation, Mechanisms, and Therapeutic Potential. Endocrinology 161. 10.1210/ENDOCR/BQZ046.

17. Kulkarni, P.P., Ekhlak, M., Singh, V., Kailashiya, V., Singh, N., and Dash, D. (2023). Fatty acid oxidation fuels agonist-induced platelet activation and thrombus formation: Targeting β-oxidation of fatty acids as an effective anti-platelet strategy. FASEB J 37. 10.1096/FJ.202201321RR.

18. Kulkarni, P.P., Ekhlak, M., and Dash, D. (2023). Energy metabolism in platelets fuels thrombus formation: Halting the thrombosis engine with small-molecule modulators of platelet metabolism. Metabolism 145, 155596. 10.1016/J.METABOL.2023.155596.

19. Öztürk, M., Freiwald, A., Cartano, J., Schmitt, R., Dejung, M., Luck, K., Al-Sady, B., Braun, S., Levin, M., and Butter, F. (2022). Proteome effects of genome-wide single gene perturbations. Nature Communications 2022 13:1 13, 1–10. 10.1038/s41467-022-33814-8.

20. Tanner, L.B., Goglia, A.G., Wei, M.H., Sehgal, T., Parsons, L.R., Park, J.O., White, E., Toettcher, J.E., and Rabinowitz, J.D. (2018). Four Key Steps Control Glycolytic Flux in Mammalian Cells. Cell Syst 7, 49-62.e8. 10.1016/J.CELS.2018.06.003.

21. Stoeger, T., Gerlach, M., Morimoto, R.I., and Nunes Amaral, L.A. (2018). Large-scale investigation of the reasons why potentially important genes are ignored. PLoS Biol 16, e2006643. 10.1371/JOURNAL.PBIO.2006643.

22. Li, H., Rukina, D., David, F.P.A., Yang Li, T., Oh, C.M., Gao, A.W., Katsyuba, E., Sleiman, M.B., Komljenovic, A., Huang, Q., et al. (2019). Identifying gene function and module connections by the integration of multispecies expression compendia. Genome Res 29, 2034–2045. 10.1101/GR.251983.119/-/DC1.

23. Superti-Furga, G., Lackner, D., Wiedmer, T., Ingles-Prieto, A., Barbosa, B., Girardi, E., Goldmann, U., Gürtl, B., Klavins, K., Klimek, C., et al. (2020). The RESOLUTE consortium: unlocking SLC transporters for drug discovery. Nature Reviews Drug Discovery 2020 19:7 19, 429–430. 10.1038/d41573-020-00056-6.

24. Langfelder, P., and Horvath, S. (2008). WGCNA: an R package for weighted correlation network analysis. BMC Bioinformatics 9, 559. 10.1186/14712105-9-559.

25. Seldin, M.M., Koplev, S., Rajbhandari, P., Vergnes, L., Rosenberg, G.M., Meng, Y., Pan, C., Phuong, T.M.N., Gharakhanian, R., Che, N., et al. (2018). A Strategy for Discovery of Endocrine Interactions with Application to Whole-Body Metabolism. Cell Metab 27, 1138-1155.e6. 10.1016/J.CMET.2018.03.015.

26. Vitart, V., Rudan, I., Hayward, C., Gray, N.K., Floyd, J., Palmer, C.N.A., Knott, S.A., Kolcic, I., Polasek, O., Graessler, J., et al. (2008). SLC2A9 is a newly identified urate transporter influencing serum urate concentration, urate excretion and gout. Nat Genet 40, 437–442. 10.1038/NG.106.

27. Dai, Y., and Lee, C.H. (2024). Transport mechanism and structural pharmacology of human urate transporter URAT1. Cell Research 2024 34:11 34, 776–787. 10.1038/s41422-024-01023-1.

28. Bahn, A., Hagos, Y., Reuter, S., Balen, D., Brzica, H., Krick, W., Burckhardt, B.C., and Sabolić, I. (2008). Identification of a New Urate and High Affinity Nicotinate Transporter, hOAT10 (SLC22A13) *. 10.1074/jbc.M800737200.

29. Ferrannini, E., and Solini, A. (2012). SGLT2 inhibition in diabetes mellitus: rationale and clinical prospects. Nature Reviews Endocrinology 2012 8:8 8, 495–502. 10.1038/nrendo.2011.243.

30. Chao, E.C., and Henry, R.R. (2010). SGLT2 inhibition — a novel strategy for diabetes treatment. Nature Reviews Drug Discovery 2010 9:7 9, 551–559. 10.1038/nrd3180.

31. Inthasot, S., Vanderhulst, J., Janssens, P., Daele, S. Van, Hoof, E. Van, Kint, C., Iconaru, L., and de Filette, J. (2024). A novel heterozygous likely pathogenic SLC5A2 variant in a diabetic patient with glucosuria and aminoaciduria. Endocrinol Diabetes Metab Case Rep 2024. 10.1530/EDM-24-0065.

32. Novikov, A., Fu, Y., Huang, W., Freeman, B., Patel, R., van Ginkel, C., Koepsell, H., Busslinger, M., Onishi, A., Nespoux, J., et al. (2019). SGLT2 inhibition and renal urate excretion: role of luminal glucose, GLUT9, and URAT1. Am J Physiol Renal Physiol 316, F173–F185. 10.1152/AJPRENAL.00462.2018.

33. Dong, M., Chen, H., Wen, S., Yuan, Y., Yang, L., Xu, D., and Zhou, L. (2023). The Mechanism of Sodium-Glucose Cotransporter-2 Inhibitors in Reducing Uric Acid in Type 2 Diabetes Mellitus. Diabetes, Metabolic Syndrome and Obesity 16, 437–445. 10.2147/DMSO.S399343.

34. Tazawa, S., Yamato, T., Fujikura, H., Hiratochi, M., Itoh, F., Tomae, M., Takemura, Y., Maruyama, H., Sugiyama, T., Wakamatsu, A., et al. (2005). SLC5A9/SGLT4, a new Na+-dependent glucose transporter, is an essential transporter for mannose, 1,5-anhydro-D-glucitol, and fructose. Life Sci 76, 1039–1050. 10.1016/J.LFS.2004.10.016.

35. Sano, R., Shinozaki, Y., and Ohta, T. (2020). Sodium–glucose cotransporters: Functional properties and pharmaceutical potential. J Diabetes Investig 11, 770. 10.1111/JDI.13255.

36. Johnson, R.J. (2023). Intestinal Hyperuricemia as a Driving Mechanism for CKD. American Journal of Kidney Diseases 81, 127–130. 10.1053/j.ajkd.2022.08.001.

37. Zhao, J.H., Stacey, D., Eriksson, N., Macdonald-Dunlop, E., Hedman, Å.K., Kalnapenkis, A., Enroth, S., Cozzetto, D., Digby-Bell, J., Marten, J., et al. (2023). Genetics of circulating inflammatory proteins identifies drivers of immune-mediated disease risk and therapeutic targets. Nature Immunology 2023 24:9 24, 1540–1551. 10.1038/s41590-023-01588-w.

38. Sun, B.B., Chiou, J., Traylor, M., Benner, C., Hsu, Y.H., Richardson, T.G., Surendran, P., Mahajan, A., Robins, C., Vasquez-Grinnell, S.G., et al. (2023). Plasma proteomic associations with genetics and health in the UK Biobank. Nature 2023 622:7982 622, 329–338. 10.1038/s41586-023-06592-6.

39. R Core Team (2021). R: A language and environment for statistical computing. R Foundation for Statistical Computing, Vienna, Austria., URL https://www.Rproject.org/.

40. Ooms, J. (2024). A Simple and Robust JSON Parser and Generator for R [R package jsonlite version 1.8.9]. CRAN: Contributed Packages. 10.32614/CRAN.PACKAGE.JSONLITE.

41. Gu, Z., Eils, R., and Schlesner, M. (2016). Complex heatmaps reveal patterns and correlations in multidimensional genomic data. Bioinformatics 32, 2847–2849. 10.1093/BIOINFORMATICS/BTW313.

42. Wickham, H. (2016). ggplot2. 10.1007/978-3-319-24277-4.

43. Sievert, C. (2021). Interactive Web-Based Data Visualization with R, Plotly, and Shiny. J R Stat Soc Ser A Stat Soc 184, 1150–1150. 10.1111/RSSA.12692.

