## Supplemental File 1 for "Synteny – a high throughput web tool to streamline causal gene prioritisation and provide insight into protein function": Synteny.R

```
#### Dependencies ####

library(colorRamp2)
library(ComplexHeatmap)
library(DT)
library(factoextra)
library(ggbeeswarm)
library(gtools)
library(Hmisc)
library(mailtoR)
library(plotly)
library(reshape2)
library(shiny)
library(shinybrowser)
library(shinyjs)
library(shinyWidgets)
library(stringr)
library(waiter)
library(withr)
library(writexl)
library(faq)

source("www/A2F_Enrichment.R")

value <- reactiveVal(0) # Set a reactive value to change each time data is saved
saveData <- function(data, name){
    eval(call("<<-", as.name(name), data)) # Link file to a standardised name
    newval <- value() + 1 # Record upload
    value(newval) # Reactivate data storage monitor
}

a2f_meta <<- read.csv("www/2025.01.08_kp_phenotypes_undefined_filtered.csv")

all_pheno_list <- list()
for(i in sort(unique(a2f_meta$Group))){
    all_pheno_list[[i]] <- as.list(subset(a2f_meta, Group == i)$ID)
}

ortho <<- read.csv("www/2025.02.02_ORTHOLOGY-ALLIANCE_TO-HUMAN.csv")

theme_harry <- function(legend.position){
    
    if(missing(legend.position)){
        legend.position = "right"
    }
    
    theme_bw() +
        theme(panel.grid = element_blank(),
              panel.border = element_blank(),
              axis.line = element_line(colour = "black"),
              axis.ticks = element_line(colour = "black"),
              axis.text = element_text(colour = "black", size = 16),
              axis.title = element_text(colour = "black", size = 18),
              plot.title = element_text(colour = "black", size = 20, hjust = 0.5),
              strip.background = element_rect(fill = "transparent", colour = "transparent"),
              strip.text = element_text(colour = "black", size = 16),
              legend.position = legend.position)
}

# Categorical scoring system

scores <<- c("Compelling (≥ 350)" = 350,
             "Extreme (≥ 100)" = 100,
             "Very strong (≥ 30)" = 30,
             "Strong (≥ 10)" = 10,
             "Moderate (≥ 3)" = 3,
             "Anecdotal (≥ 1)" = 1)

qualify <<- function(x){
    x <- as.numeric(x)
    res <- c()
    for(i in 1:length(x)){
        res <- c(res, names(which(x[i] >= scores)[1]))
    }
    return(res)
}

#### Front ####

ui <- fluidPage(theme = "style.css", 
                useShinyjs(), useSweetAlert(), use_waiter(), shinybrowser::detect(), use_mailtoR(),
                tags$head(includeHTML("www/google-analytics.html"),
                          tags$style(HTML("
                          .navbar-nav {
                          float: none !important;
                          }
                          .navbar-nav > li:nth-child(6) {
                          float: right;
                          }
                          .navbar-nav > li:nth-child(7) {
                          float: right;
                          }"))),
                
                navbarPage(title = span(strong("Synteny"),
                                        style = "color: #6060cc"), 
                           
                           id = "menu",
                           position = "fixed-top",
                           
                           tabPanel(title = "Home",
                                    br(),
                                    htmlOutput(outputId = "text_home1"),
                                    hr(),
                                    fluidRow(
                                        column(width = 6,
                                               div(img(src = "synteny_logo.png", height = "400px", width = "360px"),
                                                   style = "text-align: center;")),
                                        column(width = 6,
                                               br(),
                                               htmlOutput(outputId = "text_home2"))),
                                    # br(),
                                    hr(),
                                    fluidRow(
                                        column(width = 6,
                                               uiOutput(outputId = "start_btn"),
                                               align = "right"),
                                        column(width = 6,
                                               actionButton(inputId = "btn_tutorial", label = "Click for tutorial", icon = icon('question')),
                                               align = "left")),
                                    br()),
                           
                           tabPanel(title = "Phenotypes",
                                    br(),
                                    htmlOutput(outputId = "text_explore"),
                                    br(),
                                    tabsetPanel(
                                        tabPanel(title = "IDs", width = 12,
                                                 br(),
                                                 htmlOutput("text_phenotype_tab"),
                                                 br(),
                                                 h4(tags$b("Filter for phenotypes in:")),
                                                 actionGroupButtons(inputIds = c("tab_hmap", "tab_enrich", "tab_all"),
                                                                    labels = list(tags$span(icon("cubes-stacked"), "Heatmap"), tags$span(icon("magnifying-glass-chart"), "Enrichment"), tags$span(icon("check"), "All")),
                                                                    direction = "horizontal"),
                                                 br(),
                                                 br(),
                                                 DTOutput(outputId = "explore_table", width = '100%'),
                                                 br(),
                                                 br()),
                                        tabPanel(title = "Single genetic correlation", width = 12,
                                                 br(),
                                                 htmlOutput(outputId = "text_gcorr"),
                                                 br(),
                                                 virtualSelectInput(inputId = "pheno_gcorr", 
                                                                    label = "Select phenotype",
                                                                    choices = prepare_choices(a2f_meta, label = Phenotype, value = ID, group_by = Group),
                                                                    width = '80%', 
                                                                    search = TRUE, searchGroup = TRUE,
                                                                    updateOn = "close",
                                                                    zIndex = 3,
                                                                    multiple = FALSE),
                                                 actionButton(inputId = "load_pheno_gcorr", label = "Load data", icon = icon("bolt")),
                                                 br(),
                                                 br(),
                                                 downloadButton(outputId = "dload_gcorr_table", label = "Download table (.xlsx)", icon = icon('circle-down')),
                                                 downloadButton(outputId = "dload_gcorr_plot", label = "Download plot (.pdf)", icon = icon('camera')),
                                                 hr(),
                                                 fluidRow(
                                                     column(width = 6,
                                                            DTOutput(outputId = "gcorr_table", width = '95%')),
                                                     column(width = 6,
                                                            br(),
                                                            plotlyOutput(outputId = "gcorr_plot", width = '100%'))),
                                                 br()),
                                        tabPanel(title = "Multiple genetic correlation", width = 12,
                                                 br(),
                                                 htmlOutput(outputId = "multi_gcorr"),
                                                 br(),
                                                 virtualSelectInput(inputId = "multi_gcorr_phenos",
                                                                    label = "Select phenotypes",
                                                                    choices = prepare_choices(a2f_meta, label = Phenotype, value = ID, group_by = Group),
                                                                    width = '80%', showValueAsTags = TRUE,
                                                                    search = TRUE, searchGroup = TRUE,
                                                                    updateOn = "close",
                                                                    zIndex = 3,
                                                                    multiple = TRUE),
                                                 actionButton(inputId = "load_multi_gcorr", label = "Load data", icon = icon("bolt")),
                                                 conditionalPanel(condition = 'input.load_multi_gcorr > 0', style = "display: inline-block;",
                                                                  downloadButton(outputId = "dload_multi_gcorr_plot", label = "Download plot (.pdf)", icon = icon('camera'))),
                                                 br(),
                                                 br(),
                                                 plotOutput(outputId = "multi_gcorr_hmap"),
                                                 br())
                                    )),
                           
                           tabPanel(title = "Data",
                                    h3(tags$b("Get Started")),
                                    htmlOutput(outputId = "text_analyse"),
                                    br(),
                                    fileInput(inputId = "data_genes", label = "Upload .csv here",
                                              buttonLabel = "Browse csv...",
                                              multiple = FALSE),
                                    h4(tags$b("Or paste genes here")),
                                    actionButton(inputId = "data_manual", label = "Open dialog box"),
                                    br(),
                                    br(),
                                    uiOutput(outputId = "post_upload"),
                                    br(),
                                    uiOutput(outputId = "post_download")),
                           
                           tabPanel(title = "HuGE",
                                    h3(tags$b("On this tab")),
                                    htmlOutput(outputId = "text_huge1"),
                                    br(),
                                    downloadButton(outputId = "dload_huge", label = "Download HuGE scores (.xlsx)", icon = icon('circle-down')),
                                    br(),
                                    br(),
                                    htmlOutput(outputId = "text_huge2"),
                                    br(),
                                    tabsetPanel(
                                        tabPanel(title = "Overview",
                                                 h3(tags$b("HuGE heatmap")),
                                                 htmlOutput(outputId = "explan_overview1"),
                                                 br(),
                                                 fluidRow(column(width = 12, align = "center",
                                                                 tags$div(id = "hmap_block",
                                                                          plotOutput(outputId = "hmap_plot", width = '100%')))),
                                                 
                                                 br(),
                                                 h3(tags$b("Configure plot settings")),
                                                 br(),
                                                 fluidRow(
                                                     column(width = 4, 
                                                            sliderInput(inputId = "hmap_min", label = "Minimum score to plot", 
                                                                        min = 0, max = 350, value = 1, 
                                                                        width = '100%')),
                                                     column(width = 8, 
                                                            prettyRadioButtons(inputId = "hmap_cols", label = "HuGE score colouring",
                                                                               choiceNames = c("Discrete", "Continuous"),
                                                                               choiceValues = c("Discrete", "Continuous"),
                                                                               selected = "Continuous",
                                                                               status = "danger", bigger = FALSE, 
                                                                               shape = "round", icon = icon('check'),
                                                                               animation = 'pulse',
                                                                               inline = TRUE)
                                                     )),
                                                 br(),
                                                 prettyRadioButtons(inputId = "hmap_alpha2", label = "Phenotype layout",
                                                                    choiceNames = c("Alphabetical", "By trait group", "Clustered"),
                                                                    choiceValues = 1:3,
                                                                    selected = 3,
                                                                    status = "danger", bigger = FALSE, 
                                                                    shape = "round", icon = icon('check'),
                                                                    animation = 'pulse',
                                                                    inline = TRUE),
                                                 br(),
                                                 conditionalPanel(condition = "input.hmap_alpha2 == 2",
                                                                  prettyRadioButtons(inputId = "group_type", label = "Label phenotypes?",
                                                                                     choices = c("No", "Yes"), 
                                                                                     selected = "No",
                                                                                     status = "danger", bigger = FALSE, 
                                                                                     shape = "round", icon = icon('check'),
                                                                                     animation = 'pulse',
                                                                                     inline = TRUE),
                                                                  br()),
                                                 prettyRadioButtons(inputId = "hmap_alpha", label = "Gene layout",
                                                                    choiceNames = c("Alphabetical", "As supplied", "Clustered"),
                                                                    choiceValues = 1:3,
                                                                    selected = 3,
                                                                    status = "danger", bigger = FALSE, 
                                                                    shape = "round", icon = icon('check'),
                                                                    animation = 'pulse',
                                                                    inline = TRUE),
                                                 br(),
                                                 h3(tags$b("Only want specific phenotypes?")),
                                                 br(),
                                                 fluidRow(
                                                     column(width = 6,
                                                            virtualSelectInput(inputId = "hmap_phenos",
                                                                               label = "Select phenotypes",
                                                                               choices = c("A", "B", "C"),
                                                                               search = TRUE, searchGroup = TRUE,
                                                                               zIndex = 3,
                                                                               multiple = TRUE)),
                                                     column(width = 6,
                                                            h4(tags$b("Or plot by group")),
                                                            actionGroupButtons(inputIds = c("hmap_all", "hmap_enrich"),
                                                                               labels = list(tags$span(icon("check"), "Plot all"), tags$span(icon("magnifying-glass-chart"), "Plot enrichment tested")),
                                                                               direction = "horizontal"))),
                                                 br(),
                                                 h3(tags$b("Happy with your plot?")),
                                                 br(),
                                                 downloadButton(outputId = "dload_hmap", label = "Download heatmap (.pdf)", icon = icon('camera')),
                                                 br(),
                                                 br()),
                                        tabPanel(title = "Raw data",
                                                 br(),
                                                 textOutput(outputId = "explan_focus"),
                                                 br(),
                                                 prettyRadioButtons(inputId = "huge_table_type", label = "Select table type",
                                                                    choices = c("Phenotype", "Gene"),
                                                                    status = "danger", bigger = FALSE,
                                                                    shape = "round", icon = icon('check'),
                                                                    animation = 'pulse'),
                                                 virtualSelectInput(inputId = "huge_table_entity", 
                                                                    label = "Select phenotype",
                                                                    choices =  c("A", "B", "C"), 
                                                                    width = '80%',
                                                                    search = TRUE, searchGroup = TRUE,
                                                                    zIndex = 3,
                                                                    multiple = FALSE),
                                                 br(),
                                                 DTOutput(outputId = "huge_dt", width = '95%'),
                                                 br(),
                                                 br()),
                                        tabPanel(title = "HuGE correlations",
                                                 h3(tags$b("What is this?")),
                                                 htmlOutput("shared_text1"),
                                                 h3(tags$b("How does it work?")),
                                                 tags$ul(
                                                     tags$li("Identify clusters of genes using the heatmap."),
                                                     tags$li("Click and drag over the heatmap to select a cluster. If there are too many genes showing, use the settings below to make the plot less crowded."),
                                                     tags$li("Confirm that correct genes have been selected by checking the text box below the heatmap."),
                                                     tags$li("Scroll past the heatmap and settings to investigate if the cluster you have selected is enriched for a particular human trait."),
                                                     tags$li("Download all results by clicking the download buttons at the bottom of this page."),
                                                     tags$li("To select a different cluster of genes, double click the heatmap and then select the new cluster.")
                                                 ),
                                                 plotlyOutput(outputId = "shared_plot", width = '100%'),
                                                 br(),
                                                 verbatimTextOutput("shared_genes"),
                                                 br(),
                                                 h3(tags$b("Configure settings")),
                                                 sliderInput(inputId = "shared_min_r", label = "Set minimum correlation (R)", 
                                                             min = 0, max = 1, value = 0, 
                                                             width = '50%'),
                                                 sliderTextInput(inputId = "shared_max_p", label = "Set p value threshold", 
                                                                 choices = c(0.001, 0.01, 0.05, seq(0.1, 1, 0.1)), 
                                                                 grid = TRUE,
                                                                 selected = 1),
                                                 hr(),
                                                 h3(tags$b("Co-regulated phenotypes")),
                                                 htmlOutput("shared_text3"),
                                                 br(),
                                                 fluidRow(column(width = 3,
                                                                 numericInput(inputId = "min_shared_huge", label = "HuGE score threshold", 
                                                                              min = 1, max = 350, value = 2)),
                                                          column(width = 9,
                                                                 plotlyOutput(outputId = "shared_plot2"))),
                                                 h3(tags$b("Happy with the results?")),
                                                 br(),
                                                 downloadButton(outputId = "dload_shared_data", label = "Download correlation matrix (.xlsx)", icon = icon('circle-down')),
                                                 downloadButton(outputId = "dload_shared_plot", label = "Download correlation heatmap (.pdf)", icon = icon('camera')),
                                                 downloadButton(outputId = "dload_shared_plot2", label = "Download co-regulation plot (.pdf)", icon = icon('camera')),
                                                 br(),
                                                 br())
                                    )),
                           
                           tabPanel(title = "Enrichment",
                                    h3(tags$b("On this tab")),
                                    uiOutput(outputId = "pre_enrich"),
                                    uiOutput(outputId = "post_enrich")),
                           
                           tabPanel(title = "Help",
                                    tags$h3(tags$b("Problems?")), 
                                    tags$p("Try watching the video tutorial if you haven't already. Or take a look at our top frequently asked questions to see if any of these apply to you. If neither of these help, send us the specifics of what's not working for you along with the offending gene set and we'll get right onto resolving it for you. Click", mailtoR(email = "", text = "here", subject = "Synteny - help"), "to send us an email."),
                                    tags$h3(tags$b("Video tutorial")),
                                    HTML('<iframe width="560" height="315" src="https://www.youtube-nocookie.com/embed/eI6GQnpEpzk?si=2o4-DsZ0a_ptq5rK" title="YouTube video player" frameborder="0" allow="accelerometer; autoplay; clipboard-write; encrypted-media; gyroscope; picture-in-picture; web-share" referrerpolicy="strict-origin-when-cross-origin" allowfullscreen></iframe>'),
                                    br(), 
                                    faq(data = data.frame(question = c("What are HuGE scores? What do they mean?", 
                                                                       "What format should my data be in?", 
                                                                       "What is ortholog mapping? Why can't my genes be matched?",
                                                                       "How does human phenotype enrichment work?",
                                                                       "How do I cite Synteny?"),
                                                          answer = c(HTML("Genetic and phenotypic data are now available across large human cohorts. However, GWAS data can be difficult to interpret. To encourage greater integration of data from humans and model systems, some clever geneticists developed a simple metric to collate all of this information across studies and expand access to the insights hidden in human genetic data (Dornbos et al 2022, <a href='https://www.sciencedirect.com/science/article/pii/S1550413122001255?via%3Dihub'>Cell Metabolism</a>). The system they created is called Human Genetic Evidence, or HuGE for short.<br><br>HuGE scores provide a single interpretable measure of the strength of association between every gene and every phenotype. They are calculated using a Bayesian model to account for the effects of rare and common genetic variation, the proximity of a gene to the peak genetic association in genome wide association studies and the impact of coding mutations detected in whole exome sequencing. By aggregating genetic data across many independent studies, HuGE scores provide an unbiased and robust assessment of how strongly a gene of interest might relate to human physiology. The higher the score, the strnoger the evidence linking gene-to-phenotype."),
                                                                     HTML("Gene names should be supplied as a headerless comma-separated variable (.csv) file. Click <a href='example_genes.csv' target = 'blank', download = 'example_genes.csv'><u>here</u></a> to download an example."), 
                                                                     HTML("Orthologs are genes that are highly conserved in terms of sequence, structure and function across species. Ortholog mapping is a process to identify these equivalent genes between species. Synteny uses the <a href='https://www.alliancegenome.org/'>Alliance for Genome Resources</a> (AGR) database to perform this mapping. This allows genes from any of 8 common model organisms to be mapped to human gene names, enabling findings in model organisms to be leveraged to infer potential human disease mechanisms and gene function, and vice-versa.<br><br>This matching is only possible for official gene names. Please check problematic gene names against the AGR database to confirm that you are using the official symbols. Gene names must also be supplied as individual genes (i.e. isoform1;isoform2;isoform3 cannot be mapped, please supply as separate gene names)."),
                                                                     HTML("HPE works by comparing HuGE scores for a user-supplied gene set against those obtained by randomly sampling the genome (or a user-supplied background gene set). An average enrichment fold effect and unadjusted p value are calculated by comparing the average HuGE score of the input genes against that of the randomly sampled distribution. FDR-corrected p values are calculated by correcting for the number of phenotypes that you test enrichment for. We report both unadjusted p and FDR-corrected p values to enable HPE to be used for either targeted or exploratory analyses. If you use HPE for targeted analysis of pre-specified phenotypes, you should report both p values to enable the significance of your findings to be fully evaluated.<br><br>It is important to understand whether findings are driven by a small number of high-scoring genes or whether the signal is distributed across the input set. This can be determined by assessing where the input genes sit in relation to all other genes within the HuGE score distribution.<br><br>To determine whether HPE is biologically robust, we performed comprehensive benchmarking of its sensitivity (statistical power) and selectivity for known target traits. These analyses demonstrated that HPE can robustly detect anything over a 1.66-fold enrichment with 80% power, and that HPE consistently identified expected enrichments in curated biological pathways."),
                                                                     HTML("If Synteny has helped advance your research, please cite the preprint: Cutler et al 2025, <a href='https://www.biorxiv.org/content/10.1101/2025.03.16.643559v1.full'><u>BioRxiv</u></a."))),
                                        elementId = "faq", 
                                        faqtitle = "Frequently Asked Questions", 
                                        width = '100%')),
                           
                           tabPanel(title = "About",
                                    tags$h3(tags$b("About Us")), 
                                    tags$p("Synteny is a tool from the", a("David James Laboratory", href = "https://metabocybernetics.com/"), "at the University of Sydney, Australia. If you have suggestions for future features, click", mailtoR(email = "", text = "here", subject = "Synteny - enquiry"), "to send us an email."), 
                                    tags$h3(tags$b("Global Impact")), 
                                    tags$p("Researchers all across the globe are using Synteny to bridge the gap between large-scale human genetic studies and research in model systems. Take a look at the map below to see where our users are based. Click and drag to zoom in on a particular region."),
                                    div(htmlOutput(outputId = "user_map"),
                                        style = "text-align: center;"),
                                    br(),
                                    hr(),
                                    div(img(src = "usyd_logo.png", width = "225px"),
                                        style = "text-align: center;"),
                                    br())
                )
)


#### Back ####

server <- function(input, output, session){
    
    observeEvent(input$menu, {
        shinyjs::runjs("window.scrollTo(0, 0)")
    })
    
    id <- NULL
    hideTab(inputId = "menu", target = "Data")
    hideTab(inputId = "menu", target = "HuGE")
    hideTab(inputId = "menu", target = "Enrichment")
    
    output$text_home1 <- renderUI({
        
        url1 <- a("Association to Function", href = "https://a2f.hugeamp.org/")
        tagList("Welcome to Synteny - a systematic interface to the", url1, "(A2F) portal that leverages large-scale human genetic datasets to identify high priority genes and derive functional insights at scale.")
        
    })
    
    output$text_home2 <- renderUI({
        
        tagList("Synteny enables bulk querying of the A2F portal, streamlining the process of integrating human genetic evidence with data from model systems. Its operation is simple, requiring only a single comma-separated variable (.csv) file of gene names to be uploaded, and is compatible with genes from any of 8 common model organisms; gene names are automatically mapped to human orthologs using the", a("Alliance for Genome Resources", href = "https://www.alliancegenome.org/"), "database. All results can be exported for analysis in external software and data visualisation plots can be downloaded as high-resolution vector PDFs.", br(), br(),
                tags$i("Cite the preprint if you found Synteny helpful:"), br(), br(),
                tags$ul(tags$li("Synteny – a high throughput web tool to streamline causal gene prioritisation and provide insight into protein function.", a(tags$u("BioRxiv"), href = "https://www.biorxiv.org/content/10.1101/2025.03.16.643559v1.full")))
        )
        
    })
    
    output$text_explore <- renderUI({
        tagList("The Association to Function (A2F) Database contains an extensive collection of phenotypes. Return here if an ID needs clarification, or if you'd like to know the genetic correlation between phenotypes.")
    })
    
    output$text_phenotype_tab <- renderUI({
        tagList("Phenotype table last updated: 2025-01-08. For the most recent update, click", a(tags$u("here."), href = "https://t2d.hugeamp.org/research.html?pageid=kp_phenotypes%&portal=a2f"), br(), br(), 
                "To see more information for a specific phenotype, click on the ID link to go to the A2F portal.")
    })
    
    output$text_gcorr <- renderUI({
        tagList("Genetic correlation analysis quantifies the likelihood that two phenotypes have a shared genetic cause. The stronger the association, the more likely that the phenotypes are regulated by the same genes and biological pathways.", br(), br(), 
                "These are calculated similarly to a standard correlation coefficient, by asessing the covariance of the two genetic traits with respect to the SNP-based relatedness between individuals in the study population. Explore below to investigate A2F's genetic correlation data.")
    })
    
    output$multi_gcorr <- renderUI({
        tagList("Use this page to check the genetic correlations between multiple phenotypes. If you are unsure what a genetic correlation is, please go to the 'single genetic correlation' tab.")
    })
    
    output$text_analyse <- renderUI({
        tagList("Begin by uploading a list of genes for any organism. You can supply a headerless csv file (example ", downloadLink(outputId = "example_csv", label = tags$u("here")), "if you are unsure what this is). Or paste your gene set into the dialog box.", br(), br(), 
                "If you supply non-human gene names, they will be automatically converted using the", a("Alliance for Genome Resources", href = "https://www.alliancegenome.org/"), "ortholog database.", br(), br(), 
                "The user interface will load dynamically as you progress. If you click a button, notifications and progress bars will appear in the bottom right of this window.")
    })
    
    output$text_huge_dload <- renderUI({
        if(exists('huge')){
            
            matched <- which(!is.na(ortholog_data$Symbol2))
            
            if(length(matched) > 1){
                
                m1 <- paste0(length(matched), "/", length(genes), " genes matched to database.")
                
                if(length(matched) < length(genes)){
                    m1 <- paste0(m1, "<br><br>", paste0("The following could not be matched. Please check: <br>   - ", paste(genes[-matched], collapse = "<br>     - ")))
                }
                
                msg <- paste0("Download complete - ", m1)
                
                tagList(HTML(msg), hr(),
                        h3(tags$b("Where is my data?")),
                        tags$ul(
                            tags$li("HuGE scores: visualise human genetic evidence on the 'HuGE' tab. You can find this on the menu bar at the top of your screen"),
                            tags$li("Human phenotype enrichment: access this on the 'Enrichment' tab.")
                        ),
                        h3(tags$b("More genes to analyse?")),
                        "Restart the app using the 'Restart' button on the bottom of the 'Home' tab.",
                        br(),
                        br())
                
            } else {
                sendSweetAlert(title = "Error", text = "No genes matched to database - please check gene names and species are correct.",
                               type = "error", btn_labels = c("OK"), btn_colors = "#6060cc",
                               closeOnClickOutside = FALSE, showCloseButton = FALSE)
            }
        }
    })
    
    output$text_huge1 <- renderUI({
        tagList("Access the HuGE scores for your genes. Explore automatically generated visualisations below, or download the data directly using the download button. If you uploaded non-human genes, they will be displayed using their human names. Click", downloadLink(outputId = "ortho_download", label = tags$u("here")), "to download the ortholog mapping results if you are unsure of the human names for your genes.")
    })
    
    output$text_huge2 <- renderUI({
        tagList("Click", actionLink(inputId = "huge_help", label = tags$u("here")), "for an explanation of how to interpret HuGE scores.", br(), br(),
                "If you would like to know more about a phenotype, you can find a comprehensive list on the 'Phenotypes' tab.")
    })
    
    output$explan_overview1 <- renderUI({
        HTML("Visualise the HuGE scores for all your genes using this heatmap. If something doesn't look right, change the settings below. Download the heatmap as a PDF by clicking on the download button at the bottom of this page.")
    })
    
    output$shared_text1 <- renderUI({
        tagList("Synteny can provide insight into potential protein function by correlating HuGE scores across genes. Similar guilt-by-association strategies are implemented in tools such as WGCNA, GeneBridge and QENIE, all of which assign putative functions to genes by means of correlation of co-expression networks.")
    })
    
    output$shared_text3 <- renderUI({
        HTML("Determine which phenotypes are co-regulated by the genes you have selected. Exclude phenotypes with weak HuGE scores by increasing the HuGE score threshold. Hover your mouse over the points to see the gene name.")
    })
    
    output$explan_focus <- renderText({
        "Query results for a specific gene or phenotype. This produces a table of sorted results with links to the A2F database."
    })
    
    output$text_explan_enrich1 <- renderUI({
        tagList("Perform human phenotype enrichment (HPE). HPE is a statistical approach to determine whether input genes are more associated with a human phenotype than expected by chance. It works by comparing HuGE scores for a user-supplied gene set against those obtained by randomly sampling the genome (or a user-supplied background gene set).",
                br(),
                h3(tags$b("Configure a background")),
                "The default background for HPE is the human genome. If a different background is more relevant to your data, you can upload it below. For example, if you identified your genes through a proteomics study of muscle tissue, you should upload the rest of the muscle proteome as the background. Background genes must be from the same organism as your original gene set.", br(), br(),
                "If you supply a background, both default and background-corrected results will be shown.")
    })
    
    output$text_enrich <- renderUI({
        tagList("The bar plot shows human phenotypes that are significantly enriched in your gene set (unadjusted p < 0.05). Phenotypes remaining significant after correcting for multiple testing are highlighted in purple (FDR < 0.05). We report both unadjusted p and FDR significance to enable HPE to be used for either targeted or exploratory analyses. If you use HPE for targeted analysis of pre-specified phenotypes, you should report both p values to enable the significance of your findings to be fully evaluated.", br(), br(),
                "Adjust the plot appearance by changing the settings below. Download results by clicking on the download buttons at the bottom of this page.")
    })
    
    output$user_map <- renderUI({
        tags$iframe(src = "2025-05-28_user_map.html",
                    width = 750*1.2,
                    height = 350*1.2,
                    style = "border:0;")
    })
    
    observeEvent(input$btn_tutorial, {
        
        sendSweetAlert(title = "Tutorial", text = "We've teleported you to the help page. Click on the video to launch the tutorial.",
                       type = "info", btn_labels = c("OK"), btn_colors = "#6060cc",
                       closeOnClickOutside = FALSE, showCloseButton = FALSE)
        
        showTab(inputId = "menu", target = "Help", select = TRUE)
        
        # shinyjs::runjs("window.scrollTo(0, 1)")
        
    })
    
    pheno_tab <- reactiveValues()
    pheno_tab$df <- a2f_meta
    
    observeEvent(input$tab_hmap, {
        if(length(input$hmap_phenos) > 0){
            temp_df <- subset(a2f_meta, ID %in% input$hmap_phenos)
            pheno_tab$df <- temp_df
            
        } else {
            sendSweetAlert(title = "Caution", text = "No phenotypes selected for heatmap yet.",
                           type = "info", btn_labels = c("OK"), btn_colors = "#6060cc",
                           closeOnClickOutside = FALSE, showCloseButton = FALSE)
        }
    })
    
    observeEvent(input$tab_enrich, {
        if(length(input$enrich_phenos) > 0){
            temp_df <- subset(a2f_meta, ID %in% input$enrich_phenos)
            pheno_tab$df <- temp_df
        } else {
            sendSweetAlert(title = "Caution", text = "No phenotypes selected for enrichment yet.",
                           type = "info", btn_labels = c("OK"), btn_colors = "#6060cc",
                           closeOnClickOutside = FALSE, showCloseButton = FALSE)
        }
    })
    
    observeEvent(input$tab_all, {
        pheno_tab$df <- a2f_meta
    })
    
    output$explore_table <- renderDT({
        datatable(data = pheno_tab$df %>%
                      dplyr::select(ID, Phenotype, Group) %>%
                      mutate(Group = factor(stringr::str_to_sentence(Group)),
                             ID = paste0("<a href='https://a2f.hugeamp.org/phenotype.html?phenotype=", ID, "'>", ID, "</a>")),
                  rownames = FALSE,
                  escape = FALSE,
                  filter = "top",
                  options = list(lengthChange = FALSE))
    })
    
    observeEvent(input$load_pheno_gcorr, {
        
        if(input$pheno_gcorr == ""){
            
            sendSweetAlert(title = "Select a phenotype first",
                           type = "warning", btn_labels = c("OK"), btn_colors = "#6060cc",
                           closeOnClickOutside = FALSE, showCloseButton = FALSE)
            
        } else {
            
            waiter_show(id = "gcorr_table", html = spin_3(), color = "white")
            waiter_show(id = "gcorr_plot", html = spin_1(), color = "white")
            
            gcorr <- scrape_gCorr(pheno_ID = input$pheno_gcorr)
            saveData(data = gcorr, 
                     name = "gcorr_dat")
            
            output$gcorr_table <- renderDT({
                datatable(gcorr %>%
                              mutate(R = round(R, 2),
                                     p = formatC(p, format = "e", digits = 2)), rownames = FALSE, 
                          options = list(lengthChange = FALSE))
            })
            
            output$gcorr_plot <- renderPlotly({
                
                gcorr_plot <- ggplot(data = gcorr, aes(text = Phenotype, x = R, y = -log10(p))) +
                    geom_blank(aes(x = -R)) +
                    geom_hline(yintercept = -log10(0.05), lty = 2) +
                    geom_point() +
                    theme_harry() +
                    theme(plot.margin = unit(rep(0.5, 4), "inches")) +
                    labs(x = "Genetic Correlation",
                         y = "-log<sub>10</sub>(p)") +
                    scale_y_continuous(limits = c(0, NA))
                
                withr::with_options(list(digits = 2), ggplotly(gcorr_plot) %>% layout(title = list(text = isolate(input$pheno_gcorr))))
            })
            
            waiter_hide(id = "gcorr_table")
            waiter_hide(id = "gcorr_plot")
        }
    })
    
    
    
    
    observeEvent(input$load_multi_gcorr, {
        
        if(length(input$multi_gcorr_phenos) < 2){
            
            sendSweetAlert(title = "Select at least 2 phenotypes",
                           type = "warning", btn_labels = c("OK"), btn_colors = "#6060cc",
                           closeOnClickOutside = FALSE, showCloseButton = FALSE)
            
        } else {
            
            waiter_show(id = "multi_gcorr_hmap_wait", html = spin_3(), color = "white")
            
            id <<- showNotification(paste0("Acquiring data for ", length(input$multi_gcorr_phenos), " phenotypes"), duration = 0)
            
            withProgress(value = 0.01, {
                
                temp <- list()
                for(i in 1:length(input$multi_gcorr_phenos)){
                    
                    incProgress(1/length(input$multi_gcorr_phenos), message = paste("Phenotype", i, "of", length(input$multi_gcorr_phenos)))
                    
                    multi_gcorr_phenos <- c()
                    multi_gcorr_phenos <- scrape_gCorr(pheno_ID = input$multi_gcorr_phenos[i])
                    
                    
                    if(nrow(multi_gcorr_phenos) > 0){
                        
                        multi_gcorr_phenos <- multi_gcorr_phenos %>%
                            mutate(source = input$multi_gcorr_phenos[i],
                                   R = round(R, 2),
                                   p = formatC(p, format = "e", digits = 2), rownames = FALSE,
                                   options = list(lengthChange = FALSE))
                        
                        temp[[i]] <- multi_gcorr_phenos
                        names(temp)[[i]] <- input$multi_gcorr_phenos[i]
                        
                    } else {
                        temp[[i]] <- NA
                        names(temp)[[i]] <- input$multi_gcorr_phenos[i]
                    }
                }
                
                
                
                these_phenos <- names(temp)
                
                temp <- do.call('rbind', temp) %>%
                    subset(source %in% names(temp) & ID %in% names(temp))
                
                res <- matrix(data = NA,
                              nrow = length(these_phenos), ncol = length(these_phenos),
                              dimnames = list(these_phenos, these_phenos))
                
                for(i in 1:nrow(res)){
                    for(j in 1:ncol(res)){
                        this_dat <- subset(temp, (source == rownames(res)[i] & ID == colnames(res)[j]) | (source == colnames(res)[j] & ID == rownames(res)[i]))
                        if(nrow(this_dat) == 0){
                            next
                        } else {
                            res[i, j] <- max(this_dat$R)
                        }
                    }
                }
                
            })
            
            removeNotification(id)
            
            saveData(data = res,
                     name = "multi_gcorr_dat")
            
            output$multi_gcorr_hmap <- renderPlot({
                
                m <- max(abs(multi_gcorr_dat), na.rm = TRUE)
                myColours4 <- colorRamp2(breaks = seq(-m, m, (2*m/4)), colors = c("#eb6372", "#e89ed2", "white", "#7dc0ff", "#6060cc"))
                
                multi_gcorr_dat2 <- multi_gcorr_dat
                multi_gcorr_dat2[is.na(multi_gcorr_dat2)] <- 0
                
                Heatmap(matrix = multi_gcorr_dat2, name = "Genetic\ncorrelation",
                        cell_fun = function(j, i, x, y, width, height, fill) {
                            grid.text(sprintf("%.2f", multi_gcorr_dat[i, j]), x, y, gp = gpar(fontsize = 12))},
                        col = myColours4,
                        border = TRUE, rect_gp = gpar(col = "black", lwd = 0.2),
                        cluster_rows = TRUE, cluster_columns = TRUE,
                        row_names_side = "left", row_dend_side = "right",
                        row_names_gp = gpar(fontsize = 12), column_names_gp = gpar(fontsize = 12),
                        heatmap_legend_param = list(title_gp = gpar(fontsize = 14),
                                                    labels_gp = gpar(fontsize = 12),
                                                    border = TRUE,
                                                    tick_length = unit(1, "mm")))
                
            })
            
            waiter_hide(id = "multi_gcorr_hmap_wait")
            
        }
    })
    
    
    
    
    output$start_btn <- renderUI({
        actionButton(inputId = "btn_start", label = "Click to start", icon = icon('play'))
    })
    
    observeEvent(input$btn_start, {
        
        if(input$btn_start == 1){
            
            showTab(inputId = "menu", target = "Data", select = TRUE)
            updateActionButton(inputId = "btn_start", label = "Restart", icon = icon('repeat'))
            
        } else {
            
            waiter_show(html = spin_flower(), color = "#6060cc")
            Sys.sleep(2)
            waiter_hide()
            
            session$reload()
        }
    })
    
    
    file_count <- reactiveVal(0)
    observeEvent(input$data_genes, {
        
        file_count(file_count()+1)
        
        if(file_count() == 1){
            
            saveData(data = read.csv(input$data_genes$datapath, header = FALSE), 
                     name = "gene_dat")
            
            genes <- gene_dat %>%
                unlist() %>%
                unname()
            
            genes <- gsub(" ", "", genes)
            
            if(any(duplicated(genes))){
                genes <- genes[-which(duplicated(genes))]
            }
            
            saveData(data = genes, name = "genes")
            
            auto_species <- data.frame(species = c("Homo sapiens", unique(ortho$Gene1SpeciesName)),
                                       matches = NA)
            auto_species$matches[1] <- length(na.omit(match(genes, ortho$Gene2Symbol)))
            for(i in 2:nrow(auto_species)){
                auto_species$matches[i] <- length(na.omit(match(genes, subset(ortho, Gene1SpeciesName == auto_species$species[i])$Gene1Symbol)))
            }
            
            mouse_n <- auto_species$matches[which(auto_species$species == "Mus musculus")]
            rat_n <- auto_species$matches[which(auto_species$species == "Rattus norvegicus")]
            if(mouse_n == rat_n){
                auto_species$matches[which(auto_species$species == "Rattus norvegicus")] <- 0
            }
            
            this_species <- auto_species$species[which.max(auto_species$matches)]
            
            showNotification(paste(length(genes), " genes uploaded."), duration = 5)
            
            output$post_upload <- renderUI({
                tagList(
                    virtualSelectInput(inputId = "select_species",
                                       label = "Check the species",
                                       choices = sort(c("Homo sapiens", unique(ortho$Gene1SpeciesName))), 
                                       selected = this_species,
                                       multiple = FALSE, 
                                       width = '300px',
                                       search = FALSE),
                    h4(tags$b("Then click this button")),
                    actionButton(inputId = "btn_download", label = "Acquire HuGE scores", icon = icon('bolt'))
                )
            })
            
        } else {
            sendSweetAlert(title = "Restart required", text = "Please restart the app via the 'Home' tab if you would like to analyse another gene set.", 
                           type = "warning", btn_labels = c("OK"), btn_colors = "#6060cc",
                           closeOnClickOutside = FALSE, showCloseButton = FALSE)
        }
    })
    
    observeEvent(input$btn_download, {
        
        if(input$btn_download > 1){
            
            sendSweetAlert(title = "Good news!", text = "HuGE scores already acquired",
                           type = "info", btn_labels = c("OK"), btn_colors = "#6060cc",
                           closeOnClickOutside = FALSE, showCloseButton = FALSE)
            
        } else {
            
            # id <<- showNotification(paste("Downloading data, please wait ..."), duration = 0)
            # huge <- scrape_HugeGene_multiCore(prot_symbols = genes, max_cores = detectCores()) # Retrieve HuGE scores for selected genes
            # huge <- huge[!duplicated(huge), ] # Remove duplicates
            # removeNotification(id)
            # saveData(data = huge, name = "huge")
            
            if(input$select_species != "Homo sapiens"){
                ortho2 <- subset(ortho, Gene1SpeciesName == input$select_species)
                m <- match(genes, ortho2$Gene1Symbol)
                
                ortho_dat <- data.frame("Symbol1" = genes,
                                        "Symbol2" = ortho2[m, "Gene2Symbol"])
            } else {
                ortho_dat <- data.frame("Symbol1" = genes,
                                        "Symbol2" = ortho[match(genes, ortho$Gene2Symbol), "Gene2Symbol"])
            }
            
            saveData(data = ortho_dat, 
                     name = "ortholog_data")
            
            gene_ids <- unique(na.omit(ortholog_data$Symbol2))
            
            if(length(gene_ids) > 1){
                
                id <<- showNotification(paste0("Orthologs mapped for ", length(gene_ids), "/", nrow(ortholog_data), " genes. Acquiring data."), duration = 0)
                
                withProgress(value = 0.01, {
                    
                    temp <- list()
                    for(i in 1:length(gene_ids)){
                        incProgress(1/length(gene_ids), message = paste("Gene", i, "of", length(gene_ids)))
                        temp[[i]] <- scrape_HugeGene(gene_ids[i])
                    }
                    removeNotification(id)
                    
                    remove <- which(lapply(temp, is.null) %>% unlist())
                    if(length(remove) > 0){
                        temp <- temp[-remove]
                    }
                    
                    temp2 <- temp
                    for(i in 1:length(temp2)){
                        temp2[[i]]$gene <- colnames(temp2[[i]])[2]
                        colnames(temp2[[i]])[2] <- "HuGE"
                    }
                    
                    temp2 <- do.call('rbind', temp2)
                    
                    d <- which(duplicated(temp2))
                    if(length(d) > 0){
                        temp2 <- temp2[-d, ]
                    }
                    
                    temp3 <- dcast(data = temp2, formula = gene ~ phenotype, value.var = "HuGE") %>%
                        tibble::column_to_rownames(var = "gene") %>%
                        as.matrix() %>%
                        t()
                    
                    huge <- temp3[-which(rowSums(temp3) == ncol(temp3)), ]
                    
                    if(any(is.na(huge))){
                        huge[which(is.na(huge))] <- 0
                    }
                    
                    huge <- huge[ , unique(temp2$gene)]
                    
                })
                
            } else {
                sendSweetAlert(title = "Error", text = "No genes matched to database - please check gene names and species are correct.",
                               type = "error", btn_labels = c("OK"), btn_colors = "#6060cc",
                               closeOnClickOutside = FALSE, showCloseButton = FALSE)
            }
            
            saveData(data = huge, name = "huge")
            
            huge_pheno_mat <- data.frame(id = rownames(huge),
                                         group = str_to_sentence(a2f_meta$Group[match(rownames(huge), a2f_meta$ID)])) %>%
                arrange(id)
            
            huge_pheno_mat$group[which(is.na(huge_pheno_mat$group))] <- "Other"
            
            huge_pheno_list <- list()
            for(i in sort(unique(huge_pheno_mat$group))){
                huge_pheno_list[[i]] <- as.list(subset(huge_pheno_mat, group == i)$id)
            }
            
            no_pheno <- which(!(unname(unlist(huge_pheno_list)) %in% a2f_meta$ID))
            if(length(no_pheno) > 0){
                other_meta <- data.frame("Phenotype" = unname(unlist(huge_pheno_list))[no_pheno],
                                         "ID" = unname(unlist(huge_pheno_list))[no_pheno],
                                         "Group" = "UNKNOWN")
                a2f_meta <- rbind(a2f_meta, other_meta)
            }
            
            updateVirtualSelect(inputId = "hmap_phenos", choices = prepare_choices(subset(a2f_meta, ID %in% unname(unlist(huge_pheno_list))), label = Phenotype, value = ID, group_by = Group),
                                selected = unname(unlist(huge_pheno_list)))
            
            updateRadioButtons(inputId = "huge_table_type", selected = "Phenotype")
            updateVirtualSelect(inputId = "huge_table_entity", 
                                label = paste("Select", tolower(input$huge_table_type)),
                                choices = prepare_choices(subset(a2f_meta, ID %in% unname(unlist(huge_pheno_list))), label = Phenotype, value = ID, group_by = Group),
                                selected = unname(unlist(huge_pheno_list))[1])
            
            observeEvent(input$huge_table_type, {
                updateVirtualSelect(inputId = "huge_table_entity", label = paste("Select", tolower(input$huge_table_type)))
                
                if(input$huge_table_type == "Phenotype" & exists('huge')){
                    updateVirtualSelect(inputId = "huge_table_entity", choices = prepare_choices(subset(a2f_meta, ID %in% unname(unlist(huge_pheno_list))), label = Phenotype, value = ID, group_by = Group))
                }
                if(input$huge_table_type == "Gene" & exists('huge')){
                    updateVirtualSelect(inputId = "huge_table_entity", choices = sort(colnames(huge)))
                }
            })
            
            showTab(inputId = "menu", target = "HuGE")
            showTab(inputId = "menu", target = "Enrichment")
            sendSweetAlert(title = "Acquisition complete", 
                           type = "success", btn_labels = c("OK"), btn_colors = "#6060cc",
                           closeOnClickOutside = FALSE, showCloseButton = FALSE)
            
            output$post_download <- renderUI({
                htmlOutput(outputId = "text_huge_dload")
            })
            
            output$pre_enrich <- renderUI({
                tagList(
                    htmlOutput(outputId = "text_explan_enrich1"),
                    br(),
                    fileInput(inputId = "data_background", label = "Upload background (optional)",
                              buttonLabel = "Browse csv...",
                              multiple = FALSE),
                    h3(tags$b("Select phenotypes to test")),
                    br(),
                    actionGroupButtons(inputIds = c("enrich_all", "enrich_hmap"),
                                       labels = list(tags$span(icon("check"), "Test all"), tags$span(icon("cubes-stacked"), "Test phenotypes showing in heatmap")),
                                       direction = "horizontal"),
                    br(),
                    br(),
                    virtualSelectInput(inputId = "enrich_phenos",
                                       label = "Or select manually",
                                       choices = prepare_choices(subset(a2f_meta, ID %in% unname(unlist(huge_pheno_list))), label = Phenotype, value = ID, group_by = Group),
                                       width = '100%', showValueAsTags = TRUE,
                                       search = TRUE, searchGroup = TRUE,
                                       multiple = TRUE,
                                       disableSelectAll = TRUE),
                    h3(tags$b("Then click this button")),
                    actionButton(inputId = "btn_enrich", label = "Perform enrichment", icon = icon('magnifying-glass-chart')),
                    hr()
                )
            })
            
            observeEvent(input$data_background, {
                saveData(data = read.csv(input$data_background$datapath, header = FALSE), 
                         name = "bg_dat")
                
                bg <- bg_dat %>%
                    unlist() %>%
                    unname()
                
                if(any(duplicated(bg))){
                    bg <- bg[-which(duplicated(bg))]
                }
                
                saveData(data = bg, name = "bg")
                showNotification(paste(length(bg), "background genes accepted"), duration = 5)
                
            })
            
            observeEvent(input$enrich_all, {
                updateVirtualSelect(inputId = "enrich_phenos", selected = unname(unlist(huge_pheno_list)))
                sendSweetAlert(title = "Caution", text = "You have selected all phenotypes. Once running, this process cannot be interrupted.",
                               type = "info", btn_labels = c("OK"), btn_colors = "#6060cc",
                               closeOnClickOutside = FALSE, showCloseButton = FALSE)
            })
            
            observeEvent(input$hmap_all, {
                updateVirtualSelect(inputId = "hmap_phenos", selected = unname(unlist(huge_pheno_list)))
            })
            
            output$hmap_plot <- renderPlot({
                
                myColours <- colorRampPalette(colors = c("white", "#7dc0ff", "#6060cc", "#23234c"))
                myColours2 <- structure(c("#eb6372", "#23234c", "#6060cc", "#7dc0ff", "grey85", "white"), names = names(scores))
                
                huge_temp <- huge[rownames(huge) %in% c(input$hmap_phenos), ]
                huge2 <- log2(huge_temp)
                
                shiny::validate(
                    need(!is.null(dim(huge2)), "Insufficient genes passing filtering - please decrease the minimum HuGE score or select more phenotypes.")
                )
                keep1 <- which(apply(huge2, 2, function(x) any(x >= log2(input$hmap_min))))
                
                shiny::validate(
                    need(length(keep1) > 1, "Insufficient genes passing filtering - please decrease the minimum HuGE score or select more phenotypes.")
                )
                
                huge3 <- huge2[, keep1]
                
                keep2 <- which(apply(huge2, 1, function(x) any(x >= log2(input$hmap_min))))
                
                shiny::validate(
                    need(length(keep2) > 1, "Insufficient genes passing filtering - please decrease the minimum HuGE score or select more phenotypes.")
                )
                
                huge3 <- huge3[keep2, ]
                huge3[which(is.infinite(huge3))] <- 0
                
                lab_seq <- c(0, ceiling(max(huge3)/2)*2)
                
                huge3 <- t(huge3)
                huge3 <- huge3[ , sort(colnames(huge3))]
                
                if(input$hmap_alpha == 1){
                    huge3 <- huge3[sort(rownames(huge3)), ]
                }
                
                if(input$hmap_alpha == 3){
                    plot_clust_rows <- TRUE
                } else {
                    plot_clust_rows <- FALSE
                }
                
                if(input$hmap_alpha2 == 3){
                    plot_clust_cols <- TRUE
                } else {
                    plot_clust_cols <- FALSE
                }
                
                brks <- seq(lab_seq[1], lab_seq[2], (lab_seq[2] - lab_seq[1])/1000)
                
                if(input$hmap_alpha2 == 2){
                    
                    pheno_groups <- a2f_meta$Group[match(colnames(huge3), a2f_meta$ID)]
                    pheno_groups[which(is.na(pheno_groups))] <- "UNKNOWN"
                    
                    huge4 <- huge3
                    colnames(huge4) <- NULL
                    
                    if(input$group_type == "Yes"){
                        
                        colnames(huge4) <- colnames(huge3)
                        
                        this_plot <- Heatmap(matrix = huge4, name = "log2(HuGE)",
                                             column_split = factor(pheno_groups, levels = sort(unique(pheno_groups))), column_title = NULL, # Split by phenotype group
                                             cluster_column_slices = FALSE, 
                                             top_annotation = HeatmapAnnotation(foo = anno_block(
                                                 gp = gpar(fill = 1:length(unique(pheno_groups))),
                                                 height = unit(0.2, "cm"))),
                                             col = myColours(1000), 
                                             border = TRUE, rect_gp = gpar(col = "black", lwd = 0.2), 
                                             cluster_rows = plot_clust_rows, cluster_columns = TRUE,
                                             row_names_side = "left", row_dend_side = "right",
                                             row_names_gp = gpar(fontsize = 12), column_names_gp = gpar(fontsize = 12),
                                             heatmap_legend_param = list(title_gp = gpar(fontsize = 14),
                                                                         labels_gp = gpar(fontsize = 12),
                                                                         border = TRUE,
                                                                         tick_length = unit(1, "mm")))
                        
                        if(input$hmap_cols == "Discrete"){
                            
                            huge4 <- 2^huge4
                            huge4 <- t(apply(huge4, 1, qualify))
                            colnames(huge4) <- colnames(huge3)
                            
                            huge4 <- huge4[row_order(draw(this_plot)), ]
                            huge4 <- huge4[ , unlist(column_order(draw(this_plot)))]
                            
                            pheno_groups2 <- suppressWarnings(pheno_groups[unlist(column_order(draw(this_plot)))])
                            
                            this_plot <- Heatmap(matrix = huge4, name = "HuGE",
                                                 column_split = factor(pheno_groups2, levels = sort(unique(pheno_groups2))), column_title = NULL, # Split by phenotype group
                                                 cluster_column_slices = FALSE, 
                                                 top_annotation = HeatmapAnnotation(foo = anno_block(
                                                     gp = gpar(fill = 1:length(unique(pheno_groups))),
                                                     height = unit(0.2, "cm"))),
                                                 col = myColours2, 
                                                 border = TRUE, rect_gp = gpar(col = "black", lwd = 0.2), 
                                                 cluster_rows = plot_clust_rows, cluster_columns = TRUE,
                                                 row_names_side = "left", row_dend_side = "right",
                                                 row_names_gp = gpar(fontsize = 12), column_names_gp = gpar(fontsize = 12),
                                                 heatmap_legend_param = list(title_gp = gpar(fontsize = 14),
                                                                             labels_gp = gpar(fontsize = 12),
                                                                             border = TRUE,
                                                                             tick_length = unit(1, "mm")))
                            
                        }
                        
                        lgd = Legend(labels = paste0(1:length(unique(pheno_groups)), ": ", sort(unique(pheno_groups))),
                                     legend_gp = gpar(fill = 1:length(unique(pheno_groups))),
                                     title_gp = gpar(fontsize = 14),
                                     labels_gp = gpar(fontsize = 12),
                                     title = "Group (left to right)", 
                                     ncol = 1,
                                     title_position = "topleft")
                        
                        draw(this_plot, annotation_legend_list = lgd, padding = unit(c(2, 2, 2, 2), "mm"))
                        
                    } else {
                        
                        this_plot <- Heatmap(matrix = huge4, name = "log2(HuGE)",
                                             column_split = factor(pheno_groups, levels = sort(unique(pheno_groups))), column_title = NULL, # Split by phenotype group
                                             cluster_column_slices = FALSE, 
                                             bottom_annotation = HeatmapAnnotation(foo = anno_block(
                                                 labels = sort(unique(pheno_groups)),
                                                 labels_gp = gpar(fontsize = 12, col = 1:length(unique(pheno_groups))), 
                                                 labels_just = "right", labels_offset = 1,
                                                 labels_rot = 90, 
                                                 gp = gpar(col = "white"))),
                                             col = myColours(1000), 
                                             border = TRUE, rect_gp = gpar(col = "black", lwd = 0.2), 
                                             cluster_rows = plot_clust_rows, cluster_columns = TRUE,
                                             row_names_side = "left", row_dend_side = "right",
                                             row_names_gp = gpar(fontsize = 12), column_names_gp = gpar(fontsize = 12),
                                             heatmap_legend_param = list(title_gp = gpar(fontsize = 14),
                                                                         labels_gp = gpar(fontsize = 12),
                                                                         border = TRUE,
                                                                         tick_length = unit(1, "mm")))
                        
                        if(input$hmap_cols == "Discrete"){
                            
                            huge4 <- 2^huge4
                            huge4 <- t(apply(huge4, 1, qualify))
                            
                            huge4 <- huge4[row_order(draw(this_plot)), ]
                            huge4 <- huge4[ , unlist(column_order(draw(this_plot)))]
                            
                            pheno_groups2 <- suppressWarnings(pheno_groups[unlist(column_order(draw(this_plot)))])
                            
                            this_plot <- Heatmap(matrix = huge4, name = "HuGE",
                                                 column_split = factor(pheno_groups2, levels = sort(unique(pheno_groups2))), column_title = NULL, # Split by phenotype group
                                                 cluster_column_slices = FALSE, 
                                                 bottom_annotation = HeatmapAnnotation(foo = anno_block(
                                                     labels = sort(unique(pheno_groups2)),
                                                     labels_gp = gpar(fontsize = 12, col = 1:length(unique(pheno_groups2))), 
                                                     labels_just = "right", labels_offset = 1,
                                                     labels_rot = 90, 
                                                     gp = gpar(col = "white"))),
                                                 col = myColours2, 
                                                 border = TRUE, rect_gp = gpar(col = "black", lwd = 0.2), 
                                                 cluster_rows = plot_clust_rows, cluster_columns = TRUE,
                                                 row_names_side = "left", row_dend_side = "right",
                                                 row_names_gp = gpar(fontsize = 12), column_names_gp = gpar(fontsize = 12),
                                                 heatmap_legend_param = list(title_gp = gpar(fontsize = 14),
                                                                             labels_gp = gpar(fontsize = 12),
                                                                             border = TRUE,
                                                                             tick_length = unit(1, "mm")))
                            
                        }
                        
                        draw(this_plot, padding = unit(c(2, 2, 2, 2), "mm"))
                        
                    }
                    
                } else {
                    
                    huge4 <- huge3
                    
                    this_plot <- Heatmap(matrix = huge4, name = "log2(HuGE)",
                                         col = myColours(1000), 
                                         border = TRUE, rect_gp = gpar(col = "black", lwd = 0.2), 
                                         cluster_rows = plot_clust_rows, cluster_columns = plot_clust_cols,
                                         row_names_side = "left", row_dend_side = "right",
                                         row_names_gp = gpar(fontsize = 12), column_names_gp = gpar(fontsize = 12),
                                         heatmap_legend_param = list(title_gp = gpar(fontsize = 14),
                                                                     labels_gp = gpar(fontsize = 12),
                                                                     border = TRUE,
                                                                     tick_length = unit(1, "mm")))
                    
                    if(input$hmap_cols == "Discrete"){
                        
                        huge4 <- 2^huge4
                        huge4 <- t(apply(huge4, 1, qualify))
                        colnames(huge4) <- colnames(huge3)
                        
                        huge4 <- huge4[row_order(draw(this_plot)), ]
                        huge4 <- huge4[ , unlist(column_order(draw(this_plot)))]
                        
                        this_plot <- Heatmap(matrix = huge4, name = "HuGE",
                                             col = myColours2, 
                                             border = TRUE, rect_gp = gpar(col = "black", lwd = 0.2), 
                                             cluster_rows = plot_clust_rows, cluster_columns = plot_clust_cols,
                                             row_names_side = "left", row_dend_side = "right",
                                             row_names_gp = gpar(fontsize = 12), column_names_gp = gpar(fontsize = 12),
                                             heatmap_legend_param = list(title_gp = gpar(fontsize = 14),
                                                                         labels_gp = gpar(fontsize = 12),
                                                                         border = TRUE,
                                                                         tick_length = unit(1, "mm")))
                        
                    } 
                    
                    draw(this_plot)
                    
                }
                
            })
            
            output$huge_dt <- renderDT({
                
                shiny::validate(
                    need(input$huge_table_entity != "", paste("Please select a", tolower(input$huge_table_type)))
                )
                
                if(input$huge_table_type == "Phenotype"){
                    this_dat <- huge[input$huge_table_entity, ] %>% sort(decreasing = TRUE) %>%
                        as.data.frame() %>%
                        rownames_to_column(var = "Gene") %>%
                        rename("HuGE" = ".") %>%
                        mutate(HuGE = round(HuGE, 2))
                }
                if(input$huge_table_type == "Gene"){
                    this_dat <- huge[ , input$huge_table_entity] %>% sort(decreasing = TRUE) %>%
                        as.data.frame() %>%
                        rownames_to_column(var = "ID") %>%
                        rename("HuGE" = ".") %>%
                        mutate(HuGE = round(HuGE, 2),
                               "Phenotype" = a2f_meta$Phenotype[match(ID, a2f_meta$ID)]) %>%
                        dplyr::select("ID", "Phenotype", "HuGE")
                }
                
                this_dat <- this_dat %>%
                    mutate(log2HuGE = round(log2(HuGE), 2))
                
                
                if(input$huge_table_type == "Phenotype"){
                    this_dat <- this_dat %>%
                        mutate(Gene = paste0("<a href='https://a2f.hugeamp.org/gene.html?gene=", Gene, "'>", Gene, "</a>"))
                }
                
                if(input$huge_table_type == "Gene"){
                    this_dat <- this_dat %>%
                        mutate(ID = paste0("<a href='https://a2f.hugeamp.org/phenotype.html?phenotype=", ID, "'>", ID, "</a>"))
                }
                
                datatable(this_dat, rownames = FALSE, escape = FALSE, 
                          options = list(lengthChange = FALSE))
                
            })
            
            output$shared_plot <- renderPlotly({
                
                huge_cor <- rcorr(huge, type = "pearson")
                
                cor_r <- huge_cor$r
                cor_p <- huge_cor$P
                
                zero_sd <- which(apply(huge, 2, function(x) sd(unname(x))) == 0)
                if(length(zero_sd) > 0){
                    cor_r <- cor_r[-zero_sd, -zero_sd]
                    cor_p <- cor_p[-zero_sd, -zero_sd]
                }
                
                cor_r[which(cor_p > input$shared_max_p)] <- 0
                
                cor_r_diag <- cor_r
                diag(cor_r_diag) <- NA
                keep <- unname(which(apply(cor_r_diag, 1, function(x) any(abs(x) >= input$shared_min_r))))
                
                shiny::validate(
                    need(length(keep) > 1, "Insufficient genes passing filtering - please decrease stringency.")
                )
                cor_r <- cor_r[keep, keep]
                
                d <- dist(x = cor_r)
                hc <- hclust(d)
                ord <- hc$order[length(hc$order):1]
                
                p <- plot_ly(x = rownames(cor_r)[ord], y = colnames(cor_r)[ord], z = cor_r[ord, ord], type = "heatmap", source = "SHARED_EVENT",
                             colors = c("#610606", "#eb6372", "#e89ed2", "white", "#7dc0ff", "#6060cc", "#23234c")) %>%
                    colorbar(limits = c(-1, 1), title = "R")
                
                p %>% 
                    layout(dragmode = "select") %>%
                    event_register("plotly_brushed")
                
            })
            
            output$shared_genes <- renderPrint({
                
                huge_cor <- rcorr(huge, type = "pearson")
                
                cor_r <- huge_cor$r
                cor_p <- huge_cor$P
                
                zero_sd <- which(apply(huge, 2, function(x) sd(unname(x))) == 0)
                if(length(zero_sd) > 0){
                    cor_r <- cor_r[-zero_sd, -zero_sd]
                    cor_p <- cor_p[-zero_sd, -zero_sd]
                }
                
                cor_r[which(cor_p > input$shared_max_p)] <- 0
                
                cor_r_diag <- cor_r
                diag(cor_r_diag) <- NA
                keep <- unname(which(apply(cor_r_diag, 1, function(x) any(abs(x) >= input$shared_min_r))))
                
                shiny::validate(
                    need(length(keep) > 1, "")
                )
                cor_r <- cor_r[keep, keep]
                
                d <- dist(x = cor_r)
                hc <- hclust(d)
                ord <- hc$order[length(hc$order):1]
                
                d <- event_data("plotly_brushed", source = "SHARED_EVENT")
                if(is.null(d)){
                    "No genes selected yet - click and drag to draw a box around genes in the heatmap." 
                } else {
                    p1 <- round(d$x[1], 0) + 1
                    p2 <- round(d$x[2], 0) + 1
                    cor_ord <- cor_r[ord, ord]
                    paste(paste(c((p2 - p1) + 1, "genes selected:"), collapse = " "), paste(colnames(cor_ord)[p1:p2], collapse = ", "), collapse = " ")
                }
                
            })
            
            output$shared_plot2 <- renderPlotly({
                
                huge_cor <- rcorr(huge, type = "pearson")
                
                cor_r <- huge_cor$r
                cor_p <- huge_cor$P
                
                zero_sd <- which(apply(huge, 2, function(x) sd(unname(x))) == 0)
                if(length(zero_sd) > 0){
                    cor_r <- cor_r[-zero_sd, -zero_sd]
                    cor_p <- cor_p[-zero_sd, -zero_sd]
                }
                
                cor_r[which(cor_p > input$shared_max_p)] <- 0
                
                cor_r_diag <- cor_r
                diag(cor_r_diag) <- NA
                keep <- unname(which(apply(cor_r_diag, 1, function(x) any(abs(x) >= input$shared_min_r))))
                
                shiny::validate(
                    need(length(keep) > 1, "")
                )
                cor_r <- cor_r[keep, keep]
                
                d <- dist(x = cor_r)
                hc <- hclust(d)
                ord <- hc$order[length(hc$order):1]
                
                d <- dist(x = cor_r)
                hc <- hclust(d)
                ord <- hc$order[length(hc$order):1]
                
                dat <- event_data("plotly_brushed", source = "SHARED_EVENT")
                
                shiny::validate(
                    need(!is.null(dat), "No genes selected.")
                )
                
                p1 <- round(dat$x[1], 0) + 1
                p2 <- round(dat$x[2], 0) + 1
                cor_ord <- cor_r[ord, ord]
                these_genes <- colnames(cor_ord)[p1:p2]
                
                raw_cor <- huge[ , which(colnames(huge) %in% these_genes)] %>%
                    as.data.frame() %>%
                    rownames_to_column(var = "ID") %>%
                    mutate(Phenotype = a2f_meta$Phenotype[match(ID, a2f_meta$ID)])
                
                raw_cor$Phenotype[which(is.na(raw_cor$Phenotype))] <- raw_cor$ID[which(is.na(raw_cor$Phenotype))]
                
                raw_cor <- raw_cor[ , c(ncol(raw_cor), 2:(1+length(these_genes)))]
                
                raw_cor2 <- raw_cor
                
                keep <- which(apply(raw_cor2[ , 2:ncol(raw_cor2)], 1, function(x) length(which(x > 1))) > 1)
                raw_cor2 <- raw_cor2[keep, ]
                
                raw_mat <- as.matrix(raw_cor[ , 2:ncol(raw_cor)])
                rownames(raw_mat) <- raw_cor$Phenotype
                
                raw_mat <- apply(raw_mat, 1, as.numeric) %>%
                    t()
                colnames(raw_mat) <- colnames(raw_cor)[-1]
                
                clust_huge <- raw_mat %>%
                    melt() %>%
                    group_by(Var1) %>%
                    mutate(m = mean(log2(value)),
                           ID = a2f_meta$ID[match(Var1, a2f_meta$Phenotype)]) %>%
                    arrange(-m)
                
                ord <- unique(clust_huge$ID)
                
                this_plot <- clust_huge %>%
                    subset(m >= input$min_shared_huge) 
                
                shiny::validate(
                    need(nrow(this_plot) > 0, "Insufficient genes passing filtering - please decrease stringency.")
                )
                
                this_plot <- this_plot %>%
                    rename("HuGE" = "value") %>%
                    ggplot(aes(x = factor(ID, levels = ord), y = log2(HuGE), text = Var2)) +
                    geom_boxplot() +
                    geom_beeswarm(aes(col = Var1), size = 2) +
                    theme_harry() +
                    theme(axis.text.x = element_text(angle = 30, hjust = 1),
                          plot.margin = unit(rep(0.5, 4), "inches")) +
                    labs(y = "log2(HuGE)",
                         x = NULL) +
                    guides(col = "none") +
                    ylim(0, NA)
                
                plotly::ggplotly(p = this_plot, tooltip = "text")
                
            })
            
        }
        
    })
    
    observeEvent(input$data_manual, {
        
        file_count(file_count()+1)
        
        if(file_count() == 1){
            
            showModal(modalDialog(
                textAreaInput(inputId = "manual_geneset", label = "Enter genes (1 per line)", value =  "", 
                              height = '400px',width = '100%'),
                easyClose = FALSE, footer = actionButton(inputId = "close", label = "Save genes")
            ))
            
            observeEvent(input$close, {
                
                removeModal()
                
                genes <- unlist(strsplit(input$manual_geneset, "\n", fixed = TRUE))
                if(any(genes == "")){
                    genes <- genes[-which(genes == "")]
                }
                
                genes <- gsub(" ", "", genes)
                
                if(any(duplicated(genes))){
                    genes <- genes[-which(duplicated(genes))]
                }
                
                saveData(data = genes, name = "genes")
                
                auto_species <- data.frame(species = c("Homo sapiens", unique(ortho$Gene1SpeciesName)),
                                           matches = NA)
                auto_species$matches[1] <- length(na.omit(match(genes, ortho$Gene2Symbol)))
                for(i in 2:nrow(auto_species)){
                    auto_species$matches[i] <- length(na.omit(match(genes, subset(ortho, Gene1SpeciesName == auto_species$species[i])$Gene1Symbol)))
                }
                
                mouse_n <- auto_species$matches[which(auto_species$species == "Mus musculus")]
                rat_n <- auto_species$matches[which(auto_species$species == "Rattus norvegicus")]
                if(mouse_n == rat_n){
                    auto_species$matches[which(auto_species$species == "Rattus norvegicus")] <- 0
                }
                
                this_species <- auto_species$species[which.max(auto_species$matches)]
                
                showNotification(paste(length(genes), " genes entered."), duration = 5)
                
                output$post_upload <- renderUI({
                    tagList(
                        virtualSelectInput(inputId = "select_species",
                                           label = "Check the species",
                                           choices = sort(c("Homo sapiens", unique(ortho$Gene1SpeciesName))),
                                           selected = this_species,
                                           multiple = FALSE,
                                           width = '300px',
                                           search = FALSE),
                        h4(tags$b("Then click this button")),
                        actionButton(inputId = "btn_download", label = "Acquire HuGE scores", icon = icon('bolt'))
                    )
                })
                
            })
            
        } else {
            
            sendSweetAlert(title = "Restart required", text = "Please restart the app via the 'Home' tab if you would like to analyse another gene set.", 
                           type = "warning", btn_labels = c("OK"), btn_colors = "#6060cc",
                           closeOnClickOutside = FALSE, showCloseButton = FALSE)
            
        }
        
    })
    
    observeEvent(input$huge_help, {
        
        showModal(modalDialog(
            h3(tags$b("Interpreting HuGE scores")),
            br(),
            tags$p("HuGE scores quantify the strength of human genetic evidence linking genes to phenotypes. The following is a useful guide:"),
            br(),
            tags$ul(
                tags$li(tags$b("≥ 350"), "= compelling"),
                tags$li(tags$b("≥ 100"), "= extreme"),
                tags$li(tags$b("≥ 30"), "= very strong"),
                tags$li(tags$b("≥ 10"), "= strong"),
                tags$li(tags$b("≥ 3"), "= moderate"),
                tags$li(tags$b("≥ 1"), "= anecdotal"),
            ), 
            br(),
            tags$p("Return to the app by clicking outside this box."),
            easyClose = TRUE, footer = NULL))
        
    })
    
    observeEvent(input$hmap_enrich, {
        if(length(input$enrich_phenos) > 0){
            updateVirtualSelect(inputId = "hmap_phenos", selected = input$enrich_phenos)
            sendSweetAlert(title = "Success", text = paste(length(input$enrich_phenos), "phenotypes selected."),
                           type = "info", btn_labels = c("OK"), btn_colors = "#6060cc",
                           closeOnClickOutside = FALSE, showCloseButton = FALSE)
        } else {
            sendSweetAlert(title = "Caution", text = "No phenotypes selected for enrichment yet.",
                           type = "info", btn_labels = c("OK"), btn_colors = "#6060cc",
                           closeOnClickOutside = FALSE, showCloseButton = FALSE)
        }
    })
    
    observeEvent(input$enrich_hmap, {
        if(length(input$hmap_phenos) > 0){
            
            huge_temp <- huge[rownames(huge) %in% c(input$hmap_phenos), ]
            huge_temp[which(is.na(huge_temp))] <- 0
            huge2 <- log2(huge_temp)
            
            keep1 <- which(apply(huge2, 2, function(x) any(x >= log2(input$hmap_min))))
            huge3 <- huge2[, keep1]
            
            keep2 <- which(apply(huge2, 1, function(x) any(x >= log2(input$hmap_min))))
            huge3 <- huge3[keep2, ]
            huge3[which(is.infinite(huge3))] <- 0
            
            lab_seq <- c(0, ceiling(max(huge3)/2)*2)
            
            huge3 <- t(huge3)
            huge3 <- huge3[ , sort(colnames(huge3))]
            
            updateVirtualSelect(inputId = "enrich_phenos", selected = colnames(huge3))
            sendSweetAlert(title = "Success", text = paste(ncol(huge3), "phenotypes selected."),
                           type = "info", btn_labels = c("OK"), btn_colors = "#6060cc",
                           closeOnClickOutside = FALSE, showCloseButton = FALSE)
        } else {
            sendSweetAlert(title = "Caution", text = "No phenotypes selected for heatmap yet.",
                           type = "info", btn_labels = c("OK"), btn_colors = "#6060cc",
                           closeOnClickOutside = FALSE, showCloseButton = FALSE)
        }
    })
    
    observeEvent(input$huge_table_type, {
        if(input$huge_table_type == "Phenotype" & exists('huge')){
            updateVirtualSelect(inputId = "huge_table_entity", choices = prepare_choices(a2f_meta, label = Phenotype, value = ID, group_by = Group))
        }
        if(input$huge_table_type == "Gene" & exists('huge')){
            updateVirtualSelect(inputId = "huge_table_entity", choices = sort(colnames(huge)))
        }
    })
    
    observeEvent(input$btn_enrich, {
        
        these_phenos <- sort(input$enrich_phenos)
        
        if(length(these_phenos) >= 1){
            
            if(exists('bg')){
                
                if(input$select_species != "Homo sapiens"){
                    ortho2 <- subset(ortho, Gene1SpeciesName == input$select_species)
                    m <- match(bg, ortho2$Gene1Symbol)
                    
                    ortho_dat <- data.frame("Symbol1" = bg,
                                            "Symbol2" = ortho2[m, "Gene2Symbol"])
                } else {
                    ortho_dat <- data.frame("Symbol1" = bg,
                                            "Symbol2" = ortho[match(bg, ortho$Gene2Symbol), "Gene2Symbol"])
                }
                
                univ <- ortho_dat$Symbol2[which(!is.na(ortho_dat$Symbol2))]
                
            } else {
                univ <- NA
            }
            
            # id <<- showNotification(paste("Performing HPE, please wait ..."), duration = 0)
            # 
            # these_phenos <- sort(rownames(huge))
            # enrich <- huge_enrich(select_prots = genes,
            #                       select_phenos = these_phenos,
            #                       prot_universe = univ, # Set to NA if no background, otherwise set to vector of genes in tissue
            #                       n_perm = 10000,
            #                       huge_scores = huge)
            # removeNotification(id)
            
            id <<- showNotification(paste("Performing HPE, please stay on this page."), duration = 0)
            withProgress(message = paste("Phenotype", 1, "of", length(these_phenos)), value = 0.01, {
                
                incProgress(1/length(these_phenos), message = paste("Phenotype", 1, "of", length(these_phenos)))
                all_enrich <- huge_enrich(select_pheno = these_phenos[1], 
                                          huge_scores = huge, 
                                          prot_universe = univ, 
                                          n_perm = 10000)
                
                if(length(these_phenos) > 1){
                    for(k in 2:length(these_phenos)){
                        incProgress(1/length(these_phenos), message = paste("Phenotype", k, "of", length(these_phenos)))
                        temp <- huge_enrich(select_pheno = these_phenos[k], 
                                            huge_scores = huge, 
                                            prot_universe = univ, 
                                            n_perm = 10000)
                        
                        all_enrich$DEFAULT <- rbind(all_enrich$DEFAULT, temp$DEFAULT)
                        if(!any(is.na(univ))){ all_enrich$CORRECTED <- rbind(all_enrich$CORRECTED, temp$CORRECTED)}
                    }
                }
            })
            saveData(data = all_enrich, name = "enrich")
            removeNotification(id)
            
            sendSweetAlert(title = "Enrichment complete", text = "Scroll down to see results",
                           type = "success", btn_labels = c("OK"), btn_colors = "#6060cc",
                           closeOnClickOutside = FALSE, showCloseButton = FALSE)
            
            updatePrettyRadioButtons(inputId = "enrich_type", choiceNames = stringr::str_to_sentence(names(enrich)), choiceValues = names(enrich), prettyOptions = list(status = "danger"))
            
            output$post_enrich <- renderUI({
                tagList(
                    h3(tags$b("Explore HPE results")),
                    htmlOutput(outputId = "text_enrich"),
                    br(),
                    fluidRow(
                        column(width = 8,
                               h3(tags$b("Enriched phenotypes")),
                               br(),
                               plotOutput(outputId = "enrich_plot", width = '100%')),
                        column(width = 4,
                               h3(tags$b("Bar plot settings")),
                               br(),
                               prettyRadioButtons(inputId = "enrich_type", label = "Select background type",
                                                  choices = c("A", "B"),
                                                  status = "danger", bigger = FALSE,
                                                  animation = 'pulse'),
                               br(),
                               sliderTextInput(inputId = "enrich_filter", label = "Set significance threshold (maximum p)", 
                                               choices = c(0.001, 0.01, 0.05, seq(0.1, 1, 0.1)), 
                                               grid = TRUE,
                                               selected = 1))),
                    br(),
                    h3(tags$b("Table of results")),
                    DTOutput(outputId = "enrich_table", width = '100%'),
                    br(),
                    h3(tags$b("Gene-level contribution")),
                    tags$p("HPE aggregates enrichment signals over entire gene sets. It is important to understand whether findings are driven by a small number of high-scoring genes or whether the signal is distributed across the input set. Use this module to visualise the contributions of individual genes."),
                    br(),
                    virtualSelectInput(inputId = "labelled_distrib_pheno",
                                       label = "Select enriched phenotype",
                                       choices = prepare_choices(subset(a2f_meta, ID %in% enrich[[1]]$Phenotype), 
                                                                 label = Phenotype, value = ID),
                                       width = '50%',
                                       search = TRUE, searchGroup = FALSE,
                                       multiple = FALSE,
                                       disableSelectAll = TRUE),
                    br(),
                    plotOutput(outputId = "lablled_distrib_plot", width = '100%'),
                    br(),
                    h3(tags$b("Happy with the results?")),
                    br(),
                    downloadButton(outputId = "dload_enrichment", label = "Download enrichment plot (.pdf)", icon = icon('camera')),
                    downloadButton(outputId = "dload_enrich", label = "Download results (.xlsx)", icon = icon('circle-down')),
                    downloadButton(outputId = "dload_HPE_contrib", label = "Download contributions plot (.pdf)", icon = icon('camera')),
                    br(),
                    br()
                )
            })
            
            output$enrich_plot <- renderPlot({
                
                top_terms <- arrange(enrich[[input$enrich_type]], Enrichment) %>%
                    mutate(sig = gsub("[.]", "", sig),
                           FDR = round(p.adjust(p, "fdr"), 3),
                           FDR_sig = gtools::stars.pval(FDR),
                           FDR_sig = gsub("[.]", "", FDR_sig))
                
                top_terms <- subset(top_terms, p <= input$enrich_filter & Enrichment >= 1)
                
                ggplot(data = top_terms, aes(x = log2(Enrichment), y = factor(Phenotype, levels = Phenotype))) +
                    geom_bar(stat = "identity", aes(fill = FDR < 0.05)) +
                    geom_vline(xintercept = log2(1), lty = 2) +
                    geom_text(aes(label = sig, x = log2(Enrichment) + 0.2), size = 6) +
                    theme_harry() +
                    scale_fill_manual(values = c("FALSE" = "grey80", "TRUE" = "#6060cc")) +
                    labs(x = "HuGE log2 Fold Enrichment",
                         y = NULL)
                
            })
            
            output$enrich_table <- renderDT({
                
                this_dat <- arrange(enrich[[input$enrich_type]], -Enrichment) %>%
                    rename("ID" = "Phenotype") %>%
                    mutate(Enrichment = round(Enrichment, 2),
                           p_sig = gsub("[.]", "", sig),
                           FDR = round(p.adjust(p, "fdr"), 3),
                           FDR_sig = gtools::stars.pval(FDR),
                           FDR_sig = gsub("[.]", "", FDR_sig),
                           "Phenotype" = a2f_meta$Phenotype[match(ID, a2f_meta$ID)]) %>%
                    dplyr::select("ID", "Phenotype", "Enrichment", "p", "p_sig", "FDR", "FDR_sig")
                
                this_dat <- subset(this_dat, p <= input$enrich_filter & Enrichment >= 1) %>%
                    mutate(ID = paste0("<a href='https://a2f.hugeamp.org/phenotype.html?phenotype=", ID, "'>", ID, "</a>"))
                
                datatable(this_dat, rownames = FALSE, escape = FALSE,
                          options = list(lengthChange = FALSE))
                
            })
            
            output$lablled_distrib_plot <- renderPlot({
                
                huge_distrib <- scrape_HugePheno(phenotype = input$labelled_distrib_pheno) %>%
                    mutate(rank = 1:nrow(.))
                
                text_dat <- data.frame(x = rep(nrow(huge_distrib), 5),
                                       y = c(log2(3) + 0.35, log2(10) + 0.35, log2(30) + 0.35, log2(100) + 0.35, log2(350) + 0.35),
                                       t = c("Moderate", "Strong", "Very Strong", "Extreme", "Compelling"))
                
                huge_distrib$hit <- NA
                hits <- sample(colnames(huge), replace = FALSE)
                huge_distrib$hit[match(hits, huge_distrib$gene)] <- TRUE
                
                this_max <- max(log2(huge_distrib$huge))
                
                ggplot(data = huge_distrib, aes(x = rank, y = log2(huge))) +
                    geom_point(aes(col = log2(huge))) +
                    geom_point(data = subset(huge_distrib, hit), col = "#eb6372", size = 3) +
                    ggrepel::geom_text_repel(data = subset(huge_distrib, hit & huge >= 3), col = "#eb6372", aes(label = gene), max.overlaps = 30) +
                    geom_hline(yintercept = c(log2(3), log2(10), log2(30), log2(100), log2(350)),
                               col = "grey50", lty = 2) +
                    geom_text(data = text_dat, aes(x = x, y = y, label = t), hjust = 1, size = 8,
                              col = "grey50") +
                    theme_harry() +
                    scale_colour_gradientn(colours = c("grey85", "#7dc0ff", "#6060cc", "#23234c")) +
                    guides(col = "none") +
                    labs(x = "Gene rank",
                         y = bquote(log[2]*(HuGE)),
                         title = input$labelled_distrib_pheno) +
                    scale_y_continuous(limits = c(0, ceiling(this_max/5)*5), breaks = seq(0, ceiling(this_max/5)*5, 5))
                
            })
            
        } else {
            showNotification(paste("Please select at least 1 phenotype to enrich"), duration = 5)
        }
    })
    
    
    
    
    output$example_csv <- downloadHandler(
        filename <- function(){
            "Example gene set.csv"
        },
        content = function(file) {
            file.copy("www/example_genes.csv", file)
        }
    )
    
    output$ortho_download <- downloadHandler(
        filename <- function(){
            "Synteny_OrthologMapping.xlsx"
        },
        content = function(file) {
            this_output <- ortholog_data
            colnames(this_output) <- c("Input", "Human ortholog")
            writexl::write_xlsx(this_output, path = file, col_names = TRUE, format_headers = FALSE)
        }
    )
    
    output$dload_gcorr_table <- downloadHandler(
        filename <- function(){
            paste0("Synteny_GeneticCorrelation_data_", input$pheno_gcorr, ".xlsx")
        },
        content = function(file) {
            this_output <- list()
            
            this_output$READ_ME <- data.frame("Data" = "Genetic correlation data",
                                              "About" = paste0("This file contains details of significant genetic correlations to ", input$pheno_gcorr, ". Genetic correlations represent the degree of overlap in the genetic causes of two phenotypes. They are calculated by the Association to Function database and downloaded through Synteny. They are roughly equivalent to correlation coefficients, calculated as the covariation in two traits with respect to the genetic similarity between individuals in the study population. The data sheet contains the correlations (R) and p values."))
            this_output$data <- gcorr_dat
            
            writexl::write_xlsx(x = this_output, path = file, col_names = TRUE, format_headers = FALSE)
        }
    )
    
    output$dload_gcorr_plot <- downloadHandler(
        filename <- function(){
            paste0("Synteny_GeneticCorrelation_plot_", input$pheno_gcorr, ".pdf")
        },
        content = function(file) {
            
            ggplot(data = gcorr_dat, aes(x = R, y = -log10(p))) +
                geom_blank(aes(x = -R)) +
                geom_hline(yintercept = -log10(0.05), lty = 2) +
                geom_point() +
                theme_harry() +
                labs(x = "Genetic Correlation",
                     y = bquote(-log[10]*(p))) +
                scale_y_continuous(limits = c(0, NA))
            
            ggsave(filename = file)
            
        }
    )
    
    output$dload_multi_gcorr_plot <- downloadHandler(
        filename <- function(){
            "Synteny_MultipleGeneticCorrelation_plot.pdf"
        },
        content = function(file) {
            
            m <- max(abs(multi_gcorr_dat), na.rm = TRUE)
            myColours4 <- colorRamp2(breaks = seq(-m, m, (2*m/4)), colors = c("#eb6372", "#e89ed2", "white", "#7dc0ff", "#6060cc"))
            
            multi_gcorr_dat2 <- multi_gcorr_dat
            multi_gcorr_dat2[is.na(multi_gcorr_dat2)] <- 0
            
            this_plot <- Heatmap(matrix = multi_gcorr_dat2, name = "Genetic\ncorrelation",
                                 cell_fun = function(j, i, x, y, width, height, fill) {
                                     grid.text(sprintf("%.2f", multi_gcorr_dat[i, j]), x, y, gp = gpar(fontsize = 12))},
                                 col = myColours4,
                                 border = TRUE, rect_gp = gpar(col = "black", lwd = 0.2),
                                 cluster_rows = TRUE, cluster_columns = TRUE,
                                 row_names_side = "left", row_dend_side = "right",
                                 row_names_gp = gpar(fontsize = 12), column_names_gp = gpar(fontsize = 12),
                                 heatmap_legend_param = list(title_gp = gpar(fontsize = 14),
                                                             labels_gp = gpar(fontsize = 12),
                                                             border = TRUE,
                                                             tick_length = unit(1, "mm")))
            
            pdf(file)
            draw(this_plot)
            dev.off()
            
        }
    )
    
    output$dload_huge <- downloadHandler(
        filename <- function(){
            "Synteny_scores_data.xlsx"
        },
        content = function(file) {
            huge_save <- as.data.frame(huge) %>%
                rownames_to_column(var = "ID") %>%
                mutate("Phenotype" = a2f_meta$Phenotype[match(ID, a2f_meta$ID)])
            huge_save <- huge_save[ , c(1, ncol(huge_save), 2:(1+ncol(huge)))]
            
            this_output <- list()
            this_output$READ_ME <- data.frame("Data" = "HuGE scores",
                                              "About" = "This file contains the HuGE scores for the gene set you uploaded. Each gene (column) in the data sheet has a corresponding score to a phenotype (rows) that has been acquired from the Association to Function portal. HuGE scores quantify the strength of human genetic evidence linking genes to phenotypes and can be interpreted as HuGE ≥ 350 = compelling, ≥ 100 = extreme, ≥ 30 = very strong, ≥ 10 = strong, ≥ 3 = moderate and ≥ 1 = anecdotal. HuGE scores may have many decimal places. This is how they are calculated in the Association to Function portal, and is not relevant to their interpretation.")
            this_output$OrthologMapping <- ortholog_data
            colnames(this_output$OrthologMapping) <- c("Input", "Human ortholog")
            this_output$data <- huge_save
            
            writexl::write_xlsx(x = this_output, path = file, col_names = TRUE, format_headers = FALSE)
        }
    )
    
    output$dload_enrich <- downloadHandler(
        filename <- function(){
            "Synteny_enrichment_data.xlsx"
        },
        content = function(file) {
            
            enrich_save <- enrich
            
            if(length(enrich_save) > 1){
                for(i in 1:length(enrich_save)){
                    enrich_save[[i]] <- enrich_save[[i]] %>%
                        rename("p_sig" = "sig",
                               "ID" = "Phenotype") %>%
                        mutate(FDR = round(p.adjust(p, "fdr"), 3),
                               FDR_sig = gtools::stars.pval(FDR),
                               "Phenotype" = a2f_meta$Phenotype[match(ID, a2f_meta$ID)]) %>%
                        dplyr::select("ID", "Phenotype", "Enrichment", "p", "p_sig", "FDR", "FDR_sig")
                }
                
                this_output <- list()
                this_output$READ_ME <- data.frame("Data" = "Human phenotype enrichment",
                                                  "About" = "This file contains the results of human phenotype enrichment. This assesses whether a list of genes is more associated with human phenotypes than associated by chance. The data on the next sheets contains the fold enrichment and significance for the phenotypes you tested. One sheet contains results where the background was the human genome, and the other contains results using the custom background that you provided.")
                this_output <- list(this_output[[1]], enrich_save[[1]], enrich_save[[2]])
                names(this_output) <- c("READ_ME", "Default", "Corrected")
            } else {
                enrich_save <- enrich_save$DEFAULT %>%
                    rename("p_sig" = "sig",
                           "ID" = "Phenotype") %>%
                    mutate(FDR = round(p.adjust(p, "fdr"), 3),
                           FDR_sig = gtools::stars.pval(FDR),
                           "Phenotype" = a2f_meta$Phenotype[match(ID, a2f_meta$ID)]) %>%
                    dplyr::select("ID", "Phenotype", "Enrichment", "p", "p_sig", "FDR", "FDR_sig")
                
                this_output <- list()
                this_output$READ_ME <- data.frame("Data" = "Human phenotype enrichment",
                                                  "About" = "This file contains the results of human phenotype enrichment. This assesses whether a list of genes is more associated with human phenotypes than associated by chance. The data on the next sheet contains the fold enrichment and significance for the phenotypes you tested. You did not supply a custom background, so these results are relative to the whole human genome.")
                this_output$DEFAULT <- enrich_save
            }
            
            writexl::write_xlsx(x = this_output, path = file, col_names = TRUE, format_headers = FALSE)
        }
    )
    
    output$dload_hmap <- downloadHandler(
        filename <- function(){
            "Synteny_scores_heatmap.pdf"
        },
        content = function(file_path) {
            
            myColours <- colorRampPalette(colors = c("white", "#7dc0ff", "#6060cc", "#23234c"))
            myColours2 <- structure(c("#eb6372", "#23234c", "#6060cc", "#7dc0ff", "grey85", "white"), names = names(scores))
            
            huge_temp <- huge[rownames(huge) %in% c(input$hmap_phenos), ]
            huge_temp[which(is.na(huge_temp))] <- 0
            huge2 <- log2(huge_temp)
            
            shiny::validate(
                need(!is.null(dim(huge2)), "Insufficient genes passing filtering - please decrease the minimum HuGE score or select more phenotypes.")
            )
            keep1 <- which(apply(huge2, 2, function(x) any(x >= log2(input$hmap_min))))
            
            shiny::validate(
                need(length(keep1)>1, "Insufficient genes passing filtering - please decrease the minimum HuGE score or select more phenotypes.")
            )
            
            huge3 <- huge2[, keep1]
            
            keep2 <- which(apply(huge2, 1, function(x) any(x >= log2(input$hmap_min))))
            
            shiny::validate(
                need(length(keep2)>1, "Insufficient genes passing filtering - please decrease the minimum HuGE score or select more phenotypes.")
            )
            
            huge3 <- huge3[keep2, ]
            huge3[which(is.infinite(huge3))] <- 0
            
            lab_seq <- c(0, ceiling(max(huge3)/2)*2)
            
            huge3 <- t(huge3)
            huge3 <- huge3[ , sort(colnames(huge3))]
            
            if(input$hmap_alpha == 1){
                huge3 <- huge3[sort(rownames(huge3)), ]
            }
            
            if(input$hmap_alpha == 3){
                plot_clust_rows <- TRUE
            } else {
                plot_clust_rows <- FALSE
            }
            
            if(input$hmap_alpha2 == 3){
                plot_clust_cols <- TRUE
            } else {
                plot_clust_cols <- FALSE
            }
            
            brks <- seq(lab_seq[1], lab_seq[2], (lab_seq[2] - lab_seq[1])/1000)
            
            if(input$hmap_alpha2 == 2){
                
                pheno_groups <- a2f_meta$Group[match(colnames(huge3), a2f_meta$ID)]
                pheno_groups[which(is.na(pheno_groups))] <- "UNKNOWN"
                
                huge4 <- huge3
                colnames(huge4) <- NULL
                
                if(input$group_type == "Yes"){
                    
                    colnames(huge4) <- colnames(huge3)
                    
                    this_plot <- Heatmap(matrix = huge4, name = "log2(HuGE)",
                                         column_split = factor(pheno_groups, levels = sort(unique(pheno_groups))), column_title = NULL, # Split by phenotype group
                                         cluster_column_slices = FALSE, 
                                         top_annotation = HeatmapAnnotation(foo = anno_block(
                                             gp = gpar(fill = 1:length(unique(pheno_groups))),
                                             height = unit(0.2, "cm"))),
                                         col = myColours(1000), 
                                         border = TRUE, rect_gp = gpar(col = "black", lwd = 0.2), 
                                         cluster_rows = plot_clust_rows, cluster_columns = TRUE,
                                         row_names_side = "left", row_dend_side = "right",
                                         row_names_gp = gpar(fontsize = 12), column_names_gp = gpar(fontsize = 12),
                                         heatmap_legend_param = list(title_gp = gpar(fontsize = 14),
                                                                     labels_gp = gpar(fontsize = 12),
                                                                     border = TRUE,
                                                                     tick_length = unit(1, "mm")))
                    
                    if(input$hmap_cols == "Discrete"){
                        
                        huge4 <- 2^huge4
                        huge4 <- t(apply(huge4, 1, qualify))
                        colnames(huge4)  <- colnames(huge3)
                        
                        huge4 <- huge4[row_order(draw(this_plot)), ]
                        huge4 <- huge4[ , unlist(column_order(draw(this_plot)))]
                        
                        pheno_groups2 <- suppressWarnings(pheno_groups[unlist(column_order(draw(this_plot)))])
                        
                        this_plot <- Heatmap(matrix = huge4, name = "HuGE",
                                             column_split = factor(pheno_groups2, levels = sort(unique(pheno_groups2))), column_title = NULL, # Split by phenotype group
                                             cluster_column_slices = FALSE, 
                                             top_annotation = HeatmapAnnotation(foo = anno_block(
                                                 gp = gpar(fill = 1:length(unique(pheno_groups))),
                                                 height = unit(0.2, "cm"))),
                                             col = myColours2, 
                                             border = TRUE, rect_gp = gpar(col = "black", lwd = 0.2), 
                                             cluster_rows = plot_clust_rows, cluster_columns = TRUE,
                                             row_names_side = "left", row_dend_side = "right",
                                             row_names_gp = gpar(fontsize = 12), column_names_gp = gpar(fontsize = 12),
                                             heatmap_legend_param = list(title_gp = gpar(fontsize = 14),
                                                                         labels_gp = gpar(fontsize = 12),
                                                                         border = TRUE,
                                                                         tick_length = unit(1, "mm")))
                        
                    }
                    
                    lgd = Legend(labels = paste0(1:length(unique(pheno_groups)), ": ", sort(unique(pheno_groups))),
                                 legend_gp = gpar(fill = 1:length(unique(pheno_groups))),
                                 title_gp = gpar(fontsize = 14),
                                 labels_gp = gpar(fontsize = 12),
                                 title = "Group (left to right)", 
                                 ncol = 1,
                                 title_position = "topleft")
                    
                    pdf(file = file_path, onefile = TRUE, width = shinybrowser::get_width()/96, height = (shinybrowser::get_width()*9/16)/96)
                    draw(this_plot, annotation_legend_list = lgd, padding = unit(c(2, 2, 2, 2), "mm"))
                    dev.off()
                    
                } else {
                    
                    this_plot <- Heatmap(matrix = huge4, name = "log2(HuGE)",
                                         column_split = factor(pheno_groups, levels = sort(unique(pheno_groups))), column_title = NULL, # Split by phenotype group
                                         cluster_column_slices = FALSE, 
                                         bottom_annotation = HeatmapAnnotation(foo = anno_block(
                                             labels = sort(unique(pheno_groups)),
                                             labels_gp = gpar(fontsize = 12, col = 1:length(unique(pheno_groups))), 
                                             labels_just = "right", labels_offset = 1,
                                             labels_rot = 90, 
                                             gp = gpar(col = "white"))),
                                         col = myColours(1000), 
                                         border = TRUE, rect_gp = gpar(col = "black", lwd = 0.2), 
                                         cluster_rows = plot_clust_rows, cluster_columns = TRUE,
                                         row_names_side = "left", row_dend_side = "right",
                                         row_names_gp = gpar(fontsize = 12), column_names_gp = gpar(fontsize = 12),
                                         heatmap_legend_param = list(title_gp = gpar(fontsize = 14),
                                                                     labels_gp = gpar(fontsize = 12),
                                                                     border = TRUE,
                                                                     tick_length = unit(1, "mm")))
                    
                    if(input$hmap_cols == "Discrete"){
                        
                        huge4 <- 2^huge4
                        huge4 <- t(apply(huge4, 1, qualify))
                        
                        huge4 <- huge4[row_order(draw(this_plot)), ]
                        huge4 <- huge4[ , unlist(column_order(draw(this_plot)))]
                        
                        pheno_groups2 <- suppressWarnings(pheno_groups[unlist(column_order(draw(this_plot)))])
                        
                        this_plot <- Heatmap(matrix = huge4, name = "HuGE",
                                             column_split = factor(pheno_groups2, levels = sort(unique(pheno_groups2))), column_title = NULL, # Split by phenotype group
                                             cluster_column_slices = FALSE, 
                                             bottom_annotation = HeatmapAnnotation(foo = anno_block(
                                                 labels = sort(unique(pheno_groups2)),
                                                 labels_gp = gpar(fontsize = 12, col = 1:length(unique(pheno_groups2))), 
                                                 labels_just = "right", labels_offset = 1,
                                                 labels_rot = 90, 
                                                 gp = gpar(col = "white"))),
                                             col = myColours2, 
                                             border = TRUE, rect_gp = gpar(col = "black", lwd = 0.2), 
                                             cluster_rows = plot_clust_rows, cluster_columns = TRUE,
                                             row_names_side = "left", row_dend_side = "right",
                                             row_names_gp = gpar(fontsize = 12), column_names_gp = gpar(fontsize = 12),
                                             heatmap_legend_param = list(title_gp = gpar(fontsize = 14),
                                                                         labels_gp = gpar(fontsize = 12),
                                                                         border = TRUE,
                                                                         tick_length = unit(1, "mm")))
                        
                    }
                    
                    pdf(file = file_path, onefile = TRUE, width = shinybrowser::get_width()/96, height = (shinybrowser::get_width()*9/16)/96)
                    draw(this_plot)
                    dev.off()
                    
                }
                
            } else {
                
                huge4 <- huge3
                
                this_plot <- Heatmap(matrix = huge4, name = "log2(HuGE)",
                                     col = myColours(1000), 
                                     border = TRUE, rect_gp = gpar(col = "black", lwd = 0.2), 
                                     cluster_rows = plot_clust_rows, cluster_columns = plot_clust_cols,
                                     row_names_side = "left", row_dend_side = "right",
                                     row_names_gp = gpar(fontsize = 12), column_names_gp = gpar(fontsize = 12),
                                     heatmap_legend_param = list(title_gp = gpar(fontsize = 14),
                                                                 labels_gp = gpar(fontsize = 12),
                                                                 border = TRUE,
                                                                 tick_length = unit(1, "mm")))
                
                if(input$hmap_cols == "Discrete"){
                    
                    huge4 <- 2^huge4
                    huge4 <- t(apply(huge4, 1, qualify))
                    colnames(huge4) <- colnames(huge3)
                    
                    huge4 <- huge4[row_order(draw(this_plot)), ]
                    huge4 <- huge4[ , unlist(column_order(draw(this_plot)))]
                    
                    this_plot <- Heatmap(matrix = huge4, name = "HuGE",
                                         col = myColours2, 
                                         border = TRUE, rect_gp = gpar(col = "black", lwd = 0.2), 
                                         cluster_rows = plot_clust_rows, cluster_columns = plot_clust_cols,
                                         row_names_side = "left", row_dend_side = "right",
                                         row_names_gp = gpar(fontsize = 12), column_names_gp = gpar(fontsize = 12),
                                         heatmap_legend_param = list(title_gp = gpar(fontsize = 14),
                                                                     labels_gp = gpar(fontsize = 12),
                                                                     border = TRUE,
                                                                     tick_length = unit(1, "mm")))
                    
                } 
                
                pdf(file = file_path, onefile = TRUE, width = shinybrowser::get_width()/96, height = (shinybrowser::get_width()*9/16)/96)
                draw(this_plot)
                dev.off()
                
            }
        }
    )
    
    output$dload_shared_data <- downloadHandler(
        filename <- function(){
            "Synteny_HuGE_correlations_data.xlsx"
        },
        content = function(file) {
            
            huge_cor <- rcorr(huge, type = "pearson")
            
            cor_r <- data.frame(huge_cor$r)
            cor_p <- data.frame(huge_cor$P)
            
            zero_sd <- which(apply(huge, 2, function(x) sd(unname(x))) == 0)
            if(length(zero_sd) > 0){
                cor_r <- cor_r[-zero_sd, -zero_sd]
                cor_p <- cor_p[-zero_sd, -zero_sd]
            }
            
            cor_r <- cbind(data.frame("Gene" = colnames(cor_r)), cor_r) %>% remove_rownames()
            cor_p <- cbind(data.frame("Gene" = colnames(cor_p)), cor_p) %>% remove_rownames()
            
            huge_cor_output <- list("r" = cor_r, "p" = cor_p)
            
            this_output <- list()
            this_output$READ_ME <- data.frame("Data" = "HuGE correlation",
                                              "About" = "This file contains the correlations between HuGE scores for the genes you uploaded. One sheet contains the correlation coefficients and the other contains the p values.")
            this_output$OrthologMapping <- ortholog_data
            colnames(this_output$OrthologMapping) <- c("Input", "Human ortholog")
            
            this_output <- list(this_output[[1]], this_output[[2]], huge_cor_output[[1]], huge_cor_output[[2]])
            names(this_output) <- c("READ_ME", "OrthologMapping", "R", "p")
            
            writexl::write_xlsx(x = this_output, path = file, col_names = TRUE, format_headers = FALSE)
            
        }
    )
    
    output$dload_shared_plot <- downloadHandler(
        filename <- function(){
            "Synteny_HuGE_correlations_heatmap.pdf"
        },
        content = function(file_path) {
            
            huge_cor <- rcorr(huge, type = "pearson")
            
            cor_r <- huge_cor$r
            cor_p <- huge_cor$P
            
            zero_sd <- which(apply(huge, 2, function(x) sd(unname(x))) == 0)
            if(length(zero_sd) > 0){
                cor_r <- cor_r[-zero_sd, -zero_sd]
                cor_p <- cor_p[-zero_sd, -zero_sd]
            }
            
            cor_r[which(cor_p > input$shared_max_p)] <- 0
            
            cor_r_diag <- cor_r
            diag(cor_r_diag) <- NA
            keep <- unname(which(apply(cor_r_diag, 1, function(x) any(abs(x) >= input$shared_min_r))))
            
            shiny::validate(
                need(length(keep) > 1, "Insufficient genes passing filtering - please decrease stringency.")
            )
            cor_r <- cor_r[keep, keep]
            
            myColours3 <- colorRamp2(breaks = seq(-1, 1, (2/6)), colors = c("#610606", "#eb6372", "#e89ed2", "white", "#7dc0ff", "#6060cc", "#23234c"))
            
            this_plot <- Heatmap(matrix = cor_r, name = "R",
                                 col = myColours3, 
                                 border = TRUE, rect_gp = gpar(col = "black", lwd = 0.2), 
                                 cluster_rows = TRUE, cluster_columns = TRUE,
                                 row_names_side = "left", row_dend_side = "right",
                                 row_names_gp = gpar(fontsize = 12), column_names_gp = gpar(fontsize = 12),
                                 heatmap_legend_param = list(title_gp = gpar(fontsize = 14),
                                                             labels_gp = gpar(fontsize = 12),
                                                             border = TRUE,
                                                             tick_length = unit(1, "mm"),
                                                             at = seq(-1, 1, 0.5)))
            
            pdf(file = file_path, onefile = TRUE, width = shinybrowser::get_width()/96, height = shinybrowser::get_height()/96)
            print(this_plot)
            dev.off()
            
        }
    )
    
    output$dload_shared_plot2 <- downloadHandler(
        filename <- function(){
            "Synteny_HuGE_correlations_boxplot.pdf"
        },
        content = function(file) {
            
            huge_cor <- rcorr(huge, type = "pearson")
            
            cor_r <- huge_cor$r
            cor_p <- huge_cor$P
            
            zero_sd <- which(apply(huge, 2, function(x) sd(unname(x))) == 0)
            if(length(zero_sd) > 0){
                cor_r <- cor_r[-zero_sd, -zero_sd]
                cor_p <- cor_p[-zero_sd, -zero_sd]
            }
            
            cor_r[which(cor_p > input$shared_max_p)] <- 0
            
            cor_r_diag <- cor_r
            diag(cor_r_diag) <- NA
            keep <- unname(which(apply(cor_r_diag, 1, function(x) any(abs(x) >= input$shared_min_r))))
            
            shiny::validate(
                need(length(keep) > 1, "")
            )
            cor_r <- cor_r[keep, keep]
            
            d <- dist(x = cor_r)
            hc <- hclust(d)
            ord <- hc$order[length(hc$order):1]
            
            d <- dist(x = cor_r)
            hc <- hclust(d)
            ord <- hc$order[length(hc$order):1]
            
            dat <- event_data("plotly_brushed", source = "SHARED_EVENT")
            
            shiny::validate(
                need(!is.null(dat), "No genes selected.")
            )
            
            p1 <- round(dat$x[1], 0) + 1
            p2 <- round(dat$x[2], 0) + 1
            cor_ord <- cor_r[ord, ord]
            these_genes <- colnames(cor_ord)[p1:p2]
            
            raw_cor <- huge[ , which(colnames(huge) %in% these_genes)] %>%
                as.data.frame() %>%
                rownames_to_column(var = "ID") %>%
                mutate(Phenotype = a2f_meta$Phenotype[match(ID, a2f_meta$ID)])
            
            raw_cor$Phenotype[which(is.na(raw_cor$Phenotype))] <- raw_cor$ID[which(is.na(raw_cor$Phenotype))]
            
            raw_cor <- raw_cor[ , c(ncol(raw_cor), 2:(1+length(these_genes)))]
            
            raw_cor2 <- raw_cor
            
            keep <- which(apply(raw_cor2[ , 2:ncol(raw_cor2)], 1, function(x) length(which(x > 1))) > 1)
            raw_cor2 <- raw_cor2[keep, ]
            
            raw_mat <- as.matrix(raw_cor[ , 2:ncol(raw_cor)])
            rownames(raw_mat) <- raw_cor$Phenotype
            
            raw_mat <- apply(raw_mat, 1, as.numeric) %>%
                t()
            colnames(raw_mat) <- colnames(raw_cor)[-1]
            
            clust_huge <- raw_mat %>%
                melt() %>%
                group_by(Var1) %>%
                mutate(m = mean(log2(value)),
                       ID = a2f_meta$ID[match(Var1, a2f_meta$Phenotype)]) %>%
                arrange(-m)
            
            ord <- unique(clust_huge$ID)
            
            this_plot <- clust_huge %>%
                subset(m >= input$min_shared_huge) 
            
            shiny::validate(
                need(nrow(this_plot) > 0, "Insufficient genes passing filtering - please decrease stringency.")
            )
            
            this_plot %>%
                rename("HuGE" = "value") %>%
                ggplot(aes(x = factor(ID, levels = ord), y = log2(HuGE))) +
                geom_boxplot() +
                geom_beeswarm(aes(col = Var1), size = 2) +
                theme_harry() +
                theme(axis.text.x = element_text(angle = 30, hjust = 1),
                      plot.margin = unit(rep(0.5, 4), "inches")) +
                labs(y = "log2(HuGE)",
                     x = NULL) +
                guides(col = "none")
            
            ggsave(filename = file)
            
        }
    )
    
    output$dload_enrichment <- downloadHandler(
        filename <- function(){
            "Synteny_enrichment_plot.pdf"
        },
        content = function(file_path) {
            
            top_terms <- arrange(enrich[[input$enrich_type]], Enrichment) %>%
                mutate(sig = gsub("[.]", "", sig),
                       FDR = round(p.adjust(p, "fdr"), 3),
                       FDR_sig = gtools::stars.pval(FDR),
                       FDR_sig = gsub("[.]", "", FDR_sig))
            
            top_terms <- subset(top_terms, p <= input$enrich_filter & Enrichment >= 1)
            
            this_plot <- ggplot(data = top_terms, aes(x = log2(Enrichment), y = factor(Phenotype, levels = Phenotype))) +
                geom_bar(stat = "identity", aes(fill = FDR < 0.05)) +
                geom_vline(xintercept = log2(1), lty = 2) +
                geom_text(aes(label = sig, x = log2(Enrichment) + 0.2), size = 6) +
                theme_harry() +
                scale_fill_manual(values = c("FALSE" = "grey80", "TRUE" = "#6060cc")) +
                labs(x = "HuGE log2 Fold Enrichment",
                     y = NULL)
            
            pdf(file = file_path, onefile = TRUE)
            print(this_plot)
            dev.off()
            
        }
    )
    
    output$dload_HPE_contrib <- downloadHandler(
        filename <- function(){
            "Synteny_HPE_contribution_plot.pdf"
        },
        content = function(file_path) {
            
            huge_distrib <- scrape_HugePheno(phenotype = input$labelled_distrib_pheno) %>%
                mutate(rank = 1:nrow(.))
            
            text_dat <- data.frame(x = rep(nrow(huge_distrib), 5),
                                   y = c(log2(3) + 0.35, log2(10) + 0.35, log2(30) + 0.35, log2(100) + 0.35, log2(350) + 0.35),
                                   t = c("Moderate", "Strong", "Very Strong", "Extreme", "Compelling"))
            
            huge_distrib$hit <- NA
            hits <- sample(colnames(huge), replace = FALSE)
            huge_distrib$hit[match(hits, huge_distrib$gene)] <- TRUE
            
            this_max <- max(log2(huge_distrib$huge))
            
            this_plot <- ggplot(data = huge_distrib, aes(x = rank, y = log2(huge))) +
                geom_point(aes(col = log2(huge))) +
                geom_point(data = subset(huge_distrib, hit), col = "#eb6372", size = 3) +
                ggrepel::geom_text_repel(data = subset(huge_distrib, hit & huge >= 3), col = "#eb6372", aes(label = gene), max.overlaps = 30) +
                geom_hline(yintercept = c(log2(3), log2(10), log2(30), log2(100), log2(350)),
                           col = "grey50", lty = 2) +
                geom_text(data = text_dat, aes(x = x, y = y, label = t), hjust = 1, size = 8,
                          col = "grey50") +
                theme_harry() +
                scale_colour_gradientn(colours = c("grey85", "#7dc0ff", "#6060cc", "#23234c")) +
                guides(col = "none") +
                labs(x = "Gene rank",
                     y = bquote(log[2]*(HuGE)),
                     title = input$labelled_distrib_pheno) +
                scale_y_continuous(limits = c(0, ceiling(this_max/5)*5), breaks = seq(0, ceiling(this_max/5)*5, 5))
            
            pdf(file = file_path, onefile = TRUE)
            print(this_plot)
            dev.off()
            
        }
    )
    
    
    
    
}

shinyApp(ui = ui, server = server) # Run app
```
