## Supplemental File 1 for "Synteny – a high throughput web tool to streamline causal gene prioritisation and provide insight into protein function": A2F_Enrichment.R

```
library(org.Hs.eg.db)
library(jsonlite)
library(plyr)
library(dplyr)
library(doSNOW)
library(doParallel)
library(ggplot2)
library(tibble)
library(corrplot)

theme_harry <- function(legend.position){
  
  if(missing(legend.position)){
    legend.position = "right"
  }
  
  theme_bw() +
    theme(panel.grid = element_blank(),
          panel.border = element_blank(),
          axis.line = element_line(colour = "black"),
          axis.ticks = element_line(colour = "black"),
          axis.text = element_text(colour = "black", size = 12),
          axis.title = element_text(colour = "black", size = 14),
          plot.title = element_text(colour = "black", size = 16, hjust = 0.5),
          strip.background = element_rect(fill = "transparent", colour = "transparent"),
          strip.text = element_text(colour = "black", size = 14),
          legend.position = legend.position)
}

## Scrape phenotype associations for a set of genes
scrape_HugeGene <- function(prot_symbol){
  
  prot_symbol <- toupper(prot_symbol)
  
  out <- read_json(paste0("https://bioindex.hugeamp.org/api/bio/query/huge?q=", prot_symbol, "&fmt=row"))
  x <- data.frame(phenotype = lapply(out$data, function(x) x$phenotype) %>% unlist,
                  huge = lapply(out$data, function(x) x$huge) %>% unlist)
  
  if(nrow(x) > 0){
    colnames(x)[2] <- prot_symbol
    x
  } else {
    return(NULL)
  }
  
  return(x)
  
}

scrape_HugeGene_multiCore <- function(prot_symbols, max_cores = detectCores()-1){
  
  gene_ids <- toupper(prot_symbols)
  
  if(length(grep(";", gene_ids)) > 0){
    gene_ids <- gene_ids[-grep(";", gene_ids)]
  }
  
  cl <- makeCluster(max_cores)
  registerDoSNOW(cl)
  
  pb <- txtProgressBar(max = length(gene_ids), style = 3, width = 30)
  progress <- function(n) setTxtProgressBar(pb, n)
  opts <- list(progress = progress)
  
  t2dkp <- list()
  t2dkp <- foreach(i = 1:length(gene_ids), .packages = c('jsonlite', 'dplyr'),
                   .options.snow = opts) %dopar% {
                     
                     out <- read_json(paste0("https://bioindex.hugeamp.org/api/bio/query/huge?q=", gene_ids[i], "&fmt=row"))
                     x <- data.frame(phenotype = lapply(out$data, function(x) x$phenotype) %>% unlist,
                                     huge = lapply(out$data, function(x) x$huge) %>% unlist)
                     
                     if(nrow(x) > 0){
                       colnames(x)[2] <- gene_ids[i]
                       x
                     } else {
                       return(NULL)
                     }
                   }
  close(pb)
  stopCluster(cl)
  
  if(any(lapply(t2dkp, is.null) %>% unlist)){
    t2dkp <- t2dkp[-which(lapply(t2dkp, is.null) %>% unlist)]
  }
  
  t2dkp <- plyr::join_all(t2dkp, by = 'phenotype', type = 'left')
  
  t2dkp <- matrix(data = unlist(t2dkp[ , -1]), 
                  nrow = nrow(t2dkp), ncol = ncol(t2dkp)-1,
                  dimnames = list(t2dkp$phenotype, colnames(t2dkp)[-1]))
  
  return(t2dkp)
  
}

## Scrape gene associations for a given phenotype
scrape_HugePheno <- function(phenotype){
  
  out <- read_json(paste0("https://bioindex.hugeamp.org/api/bio/query/huge-phenotype?q=", phenotype, "&fmt=row&limit=100000"))
  huge_pheno <- data.frame(gene = lapply(out$data, function(x) x$gene) %>% unlist,
                           huge = lapply(out$data, function(x) x$huge) %>% unlist)
  return(huge_pheno)
  
}

## Test enrichment ##
huge_enrich <- function(select_pheno, huge_scores, prot_universe, n_perm){
  
  all_enrich <- list()
  
  huge_res <- list()
  for(i in 1:length(select_pheno)){
    
    gene_scores <- data.frame(gene = names(huge_scores[select_pheno[i], ]), 
                              huge = unname(huge_scores[select_pheno[i], ])) # Reformat scores for genes
    
    pheno <- scrape_HugePheno(phenotype = select_pheno[i]) %>%
      subset(!(gene %in% gene_scores$gene)) # Get background scores for all genes
    
    # NAIVE UNIVERSE #
    
    perm <- c()
    for(j in 1:n_perm){
      perm <- c(perm, mean(pheno$huge[sample(1:nrow(pheno), nrow(gene_scores))]))
    }
    
    enrich_fold <- mean(gene_scores$huge)/mean(perm)
    enrich_p <- length(which(perm > mean(gene_scores$huge)))/length(perm)
    
    # BG_CORRECTED UNIVERSE #
    
    if(!any(is.na(prot_universe))){
      
      pheno2 <- pheno %>%
        subset(gene %in% toupper(prot_universe))
      
      perm2 <- c()
      for(j in 1:n_perm){
        perm2 <- c(perm2, mean(pheno2$huge[sample(1:nrow(pheno2), nrow(gene_scores))]))
      }
      
      enrich_fold2 <- mean(gene_scores$huge)/mean(perm2)
      enrich_p2 <- length(which(perm2 > mean(gene_scores$huge)))/length(perm2)
      
      huge_res[[select_pheno[i]]]$DEFAULT <- c(enrich_fold, enrich_p)
      huge_res[[select_pheno[i]]]$CORRECTED <- c(enrich_fold2, enrich_p2)
      
    } else {
      huge_res[[select_pheno[i]]]$DEFAULT <- c(enrich_fold, enrich_p)
    }
    
  }
  
  DEFAULT <- lapply(huge_res, function(x) x$DEFAULT)
  CORRECTED <- lapply(huge_res, function(x) x$CORRECTED)
  
  DEFAULT <- DEFAULT %>%
    unlist() %>%
    matrix(nrow = length(select_pheno), ncol = 2, byrow = TRUE, dimnames = list(select_pheno, c("Enrichment", "p"))) %>%
    as.data.frame() %>%
    mutate(sig = gtools::stars.pval(p)) %>%
    rownames_to_column(var = "Phenotype") %>%
    arrange(Enrichment)
  
  if(!any(is.na(prot_universe))){
    CORRECTED <- CORRECTED %>%
      unlist() %>%
      matrix(nrow = length(select_pheno), ncol = 2, byrow = TRUE, dimnames = list(select_pheno, c("Enrichment", "p"))) %>%
      as.data.frame() %>%
      mutate(sig = gtools::stars.pval(p)) %>%
      rownames_to_column(var = "Phenotype") %>%
      arrange(Enrichment)
    
    all_enrich$DEFAULT <- DEFAULT
    all_enrich$CORRECTED <- CORRECTED
  } else {
    all_enrich$DEFAULT <- DEFAULT
  }
  
  return(all_enrich)
  
}

huge_enrich_multiCore <- function(select_phenos, huge_scores, prot_universe, n_perm, max_cores = detectCores()){
  
  cl <- makeCluster(detectCores()-1)
  registerDoSNOW(cl)
  
  pb <- txtProgressBar(max = length(select_phenos), style = 3, width = 30)
  progress <- function(n) setTxtProgressBar(pb, n)
  opts <- list(progress = progress)
  
  huge_res <- list()
  huge_res <- foreach(this_pheno = select_phenos, .export = c('scrape_HugePheno'),
                      .packages = c('dplyr', 'jsonlite'), .options.snow = opts) %dopar% {
                        
                        gene_scores <- data.frame(gene = names(huge_scores[this_pheno, ]), 
                                                  huge = unname(huge_scores[this_pheno, ])) # Reformat scores for genes
                        
                        pheno <- scrape_HugePheno(phenotype = this_pheno) %>%
                          subset(!(gene %in% gene_scores$gene)) # Get background scores for all genes
                        
                        # NAIVE UNIVERSE #
                        
                        perm <- c()
                        for(i in 1:n_perm){
                          perm <- c(perm, mean(pheno$huge[sample(1:nrow(pheno), nrow(gene_scores))]))
                        }
                        
                        enrich_fold <- mean(gene_scores$huge)/mean(perm)
                        enrich_p <- length(which(perm > mean(gene_scores$huge)))/length(perm)
                        
                        # BG_CORRECTED UNIVERSE #
                        
                        if(!any(is.na(prot_universe))){
                          
                          pheno2 <- pheno %>%
                            subset(gene %in% toupper(prot_universe))
                          
                          perm2 <- c()
                          for(i in 1:n_perm){
                            perm2 <- c(perm2, mean(pheno2$huge[sample(1:nrow(pheno2), nrow(gene_scores))]))
                          }
                          
                          enrich_fold2 <- mean(gene_scores$huge)/mean(perm2)
                          enrich_p2 <- length(which(perm2 > mean(gene_scores$huge)))/length(perm2)
                          
                          return(list("DEFAULT" = c(enrich_fold, enrich_p),
                                      "CORRECTED" = c(enrich_fold2, enrich_p2)))
                          
                        } else {
                          return(list("DEFAULT" = c(enrich_fold, enrich_p)))
                        }
                        
                      }
  
  close(pb)
  stopCluster(cl)
  
  DEFAULT <- lapply(huge_res, function(x) x$DEFAULT)
  CORRECTED <- lapply(huge_res, function(x) x$CORRECTED)
  
  DEFAULT <- DEFAULT %>%
    unlist() %>%
    matrix(nrow = length(select_phenos), ncol = 2, byrow = TRUE, dimnames = list(select_phenos, c("Enrichment", "p"))) %>%
    as.data.frame() %>%
    mutate(sig = gtools::stars.pval(p)) %>%
    rownames_to_column(var = "Phenotype") %>%
    arrange(Enrichment)
  
  if(!any(is.na(prot_universe))){
    CORRECTED <- CORRECTED %>%
      unlist() %>%
      matrix(nrow = length(select_phenos), ncol = 2, byrow = TRUE, dimnames = list(select_phenos, c("Enrichment", "p"))) %>%
      as.data.frame() %>%
      mutate(sig = gtools::stars.pval(p)) %>%
      rownames_to_column(var = "Phenotype") %>%
      arrange(Enrichment)
    
    return(list("DEFAULT" = DEFAULT, "CORRECTED" = CORRECTED))
  } else {
    return(list("DEFAULT" = DEFAULT))
  }
  
}

## Scrape genetic correlation data ##
scrape_gCorr <- function(pheno_ID){
  
  out <- read_json(paste0("https://bioindex.hugeamp.org/api/bio/query/genetic-correlation?q=", pheno_ID, "&fmt=row&limit=100000"))
  
  gcorr <- data.frame(phenotype = lapply(out$data, function(x) x$other_phenotype) %>% unlist,
                      R = lapply(out$data, function(x) x$rg) %>% unlist,
                      p = lapply(out$data, function(x) x$pValue) %>% unlist) 
  
  if(nrow(gcorr) == 0){
    return(gcorr)
  } else {
    
    gcorr <- gcorr %>% rename("ID" = "phenotype") %>%
      mutate("Phenotype" = a2f_meta$Phenotype[match(ID, a2f_meta$ID)]) %>%
      dplyr::select(ID, Phenotype, R, p)
    
    gcorr$Phenotype[which(is.na(gcorr$Phenotype))] <- gcorr$ID[which(is.na(gcorr$Phenotype))]
    
    return(gcorr)
  }
}


## Load HuGE matrix #
format_huge <- function(x){
  matrix(unlist(x[ , -1]), ncol = ncol(x) - 1,
         dimnames = list(unname(unlist(x[ , 1])), colnames(x)[-1]))
}

## Load enrichment results ##
load_enrichment <- function(path){
  t1 <- readxl::read_xlsx(path, sheet = 1)
  t2 <- readxl::read_xlsx(path, sheet = 2)
  return(list("DEFAULT" = t1, "CORRECTED" = t2))
}
```
