## Supplementary figures and images for "Synteny – a high throughput web tool to streamline causal gene prioritisation and provide insight into protein function"

### synteny_logo.png

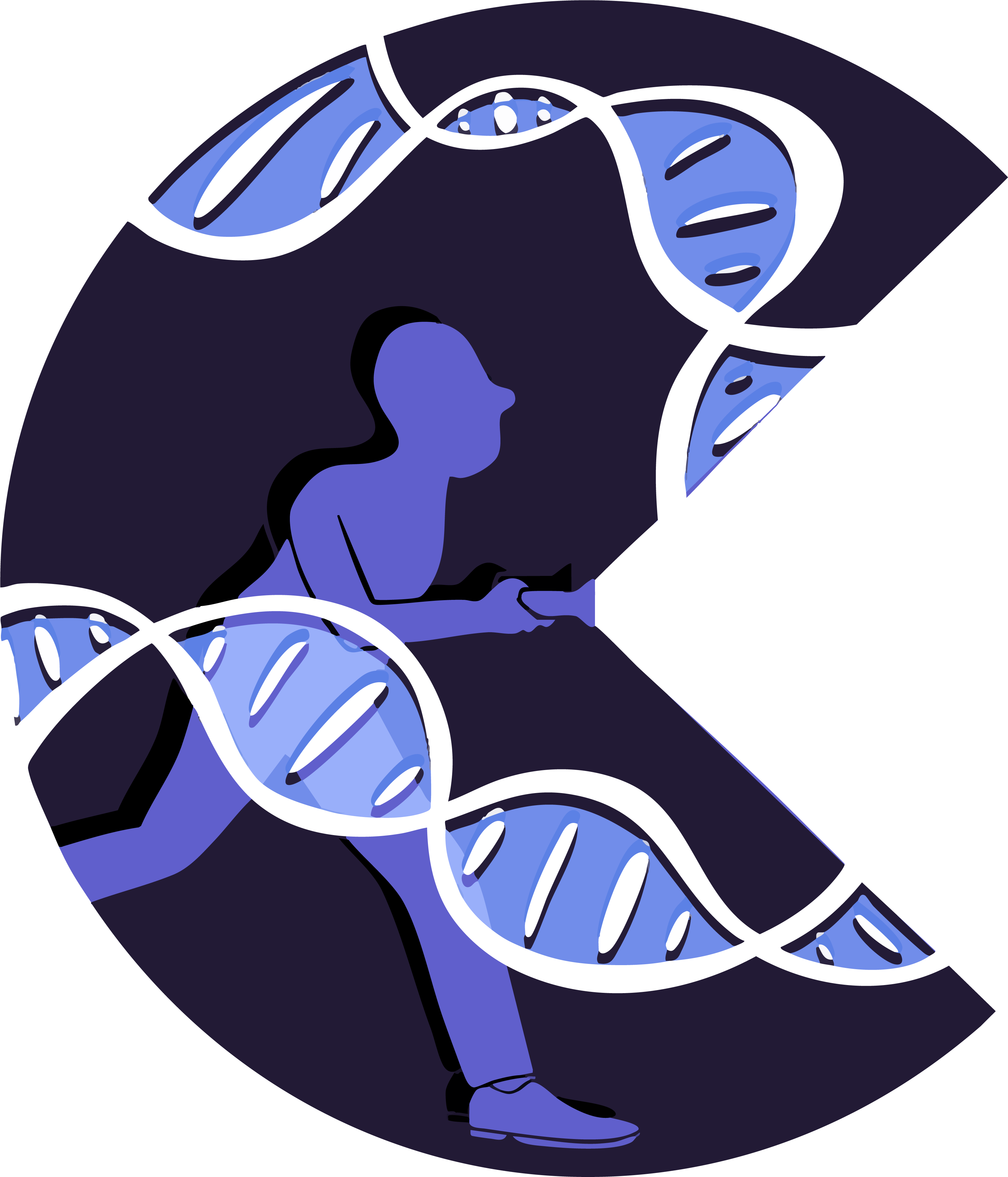

### usyd_logo.png

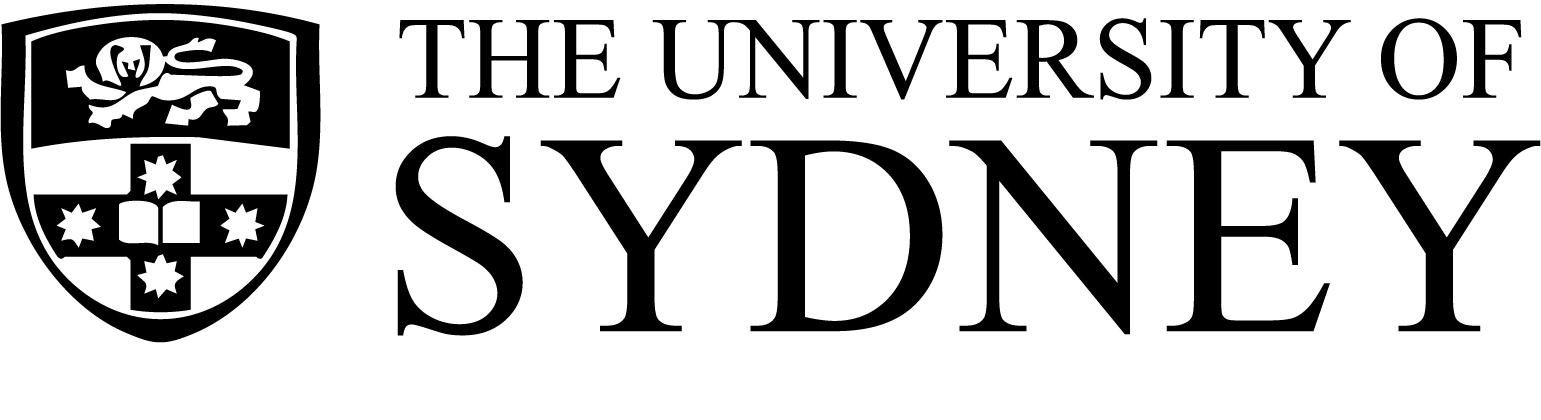
